# Proximity-assisted photoactivation (PAPA): Detecting molecular interactions in live-cell single-molecule imaging

**DOI:** 10.1101/2021.12.13.472508

**Authors:** Thomas G.W. Graham, John J. Ferrie, Gina M. Dailey, Robert Tjian, Xavier Darzacq

## Abstract

Single-molecule imaging provides a powerful way to study biochemical processes in live cells, yet it remains challenging to track single molecules while simultaneously detecting their interactions. Here we describe a novel property of rhodamine dyes, proximity-assisted photoactivation (PAPA), in which one fluorophore (the “sender”) can reactivate a second fluorophore (the “receiver”) from a dark state. PAPA requires proximity between the two fluorophores, yet it operates at a longer average intermolecular distance than Förster resonance energy transfer (FRET). We show that PAPA can be used in live cells both to detect protein-protein interactions and to highlight a sub-population of labeled protein complexes in which two different labels are in proximity. In proof-of-concept experiments, PAPA detected the expected correlation between androgen receptor self-association and chromatin binding at the single-cell level. These results establish a new way in which a photophysical property of fluorophores can be harnessed to study molecular interactions in single-molecule imaging of live cells.

## Introduction

Most proteins function by interacting with other proteins, yet we lack tools to study these potentially transient interactions at single-molecule resolution in live cells. Single particle tracking (SPT) provides a valuable tool for monitoring the motions of individual protein molecules^1–4^, but it does not distinguish compositionally and functionally distinct complexes of the same protein. Although in principle one could label two different proteins with different fluorophores and track both simultaneously, the requirement for sparse labeling of both proteins makes detection of double-labeled complexes exceedingly inefficient. Other approaches also have limitations. Single-molecule Förster resonance energy transfer (smFRET), though powerful for monitoring interactions *in vitro*, has proven technically challenging in cells due to the requirement for sparse double-labeling, the large size of genetically encoded tags relative to the working distance of FRET, and the brief observation time (tens of milliseconds) for fast-diffusing complexes^5^. Fluorescence cross-correlation spectroscopy (FCCS) can detect bulk molecular interactions, yet it does not provide spatial trajectories for individual molecules, which are useful for measuring such properties as chromatin residence time and anomalous diffusion^4,6–8^.

An alternative *in vitro* proximity sensor to smFRET was devised by Bates, Blosser, and Zhuang, who observed that exciting one cyanine dye can reactivate a nearby cyanine dye from a dark state^9^. Although photoswitching of cyanine dye pairs enabled early implementations of STORM imaging^10^, its application as a proximity sensor has been limited by the short inter-fluorophore distance required (≤ 2 nm), the poor cell permeability of cyanine dyes, and the need for high thiol concentrations and an oxygen scavenging system^11,12^.

To our knowledge, it has not been reported whether a similar process of reactivation can occur for pairs of non-cyanine dyes. However, many fluorophores—notably rhodamine dyes—can enter a dark state and be directly reactivated by short-wavelength (e.g., 405 nm) light^13^. This phenomenon, which has been employed for direct STORM (dSTORM) imaging in both live and fixed cells^14–16^, is thought to involve conversion of excited triplet state fluorophores to reduced species whose absorbance is shifted to shorter wavelengths^9,13,14,17–19^.

The development of bright, cell permeable Janelia Fluor (JF) dyes, based on rhodamine and silicon-rhodamine chemical scaffolds, has transformed single-molecule imaging in live cells^15^. Here, we show that Janelia Fluor X 650 (JFX650)^20^ and similar fluorophores can be reactivated from a dark state by excitation of a nearby fluorophore such as Janelia Fluor 549 (JF549), a phenomenon which we term **proximity-assisted photoactivation (PAPA)**. In contrast to cyanine dye reactivation, PAPA of JF dyes occurs under physiological conditions in live cells and requires neither an oxygen scavenging system nor exogenous thiols. While PAPA requires proximity between the two fluorophores, its effective distance range extends beyond that of FRET, making it a potentially more versatile interaction sensor.

Most importantly, PAPA provides a new way to detect protein interactions in live cells at single-molecule resolution. We show that PAPA can be used to detect the formation of protein dimers and that it can selectively highlight sub-populations of molecules within defined mixtures. As a further proof of concept, we combined single-molecule tracking with PAPA to analyze the increase in chromatin binding induced by self-association of androgen receptor. By enabling the previously elusive detection of protein-protein interactions, PAPA will provide a new dimension of information for live cell single-molecule imaging.

## Results

### Proximity-assisted photoactivation (PAPA) of Janelia Fluor dyes

We fortuitously discovered PAPA while imaging an oligomeric protein labeled with two different JF dyes. U2OS cells expressing Halo-tagged NPM1 (a pentameric nucleolar protein)^2^ were labeled with a low concentration of Janelia Fluor X 650 Halo-tag ligand (JFX650-HTL)^20^ to track single molecules, together with a higher concentration of Janelia Fluor 549 HaloTag ligand (JF549-HTL) to visualize nucleoli. When we alternately excited JFX650 with red light (633 nm) and JF549 with green light (561 nm), we noticed that some JFX650 molecules that had gone dark during red illumination suddenly reappeared after a brief, 7-millisecond pulse of green light (green vertical lines in Fig. 1a(i) and green box in Fig. 1b). Consistent with previous work^15^, we also observed reactivation of JFX650 by violet light, both with and without JF549-HTL (violet vertical lines in Fig. 1a(i,ii) and violet box in Fig. 1b). However, reactivation of JFX650 by green light required co-labeling with JF549 (compare Fig. 1a(i) and (ii)), implying that reactivation results not from direct absorption of green light by dark-state JFX650 but indirectly due to excitation of JF549. Green illumination of cells labeled with JF549-HTL alone did not produce localizations in the JFX650 channel, demonstrating that this effect is not due to JF549 photochromism (Fig. 1a(iii)).

**Figure 1:**
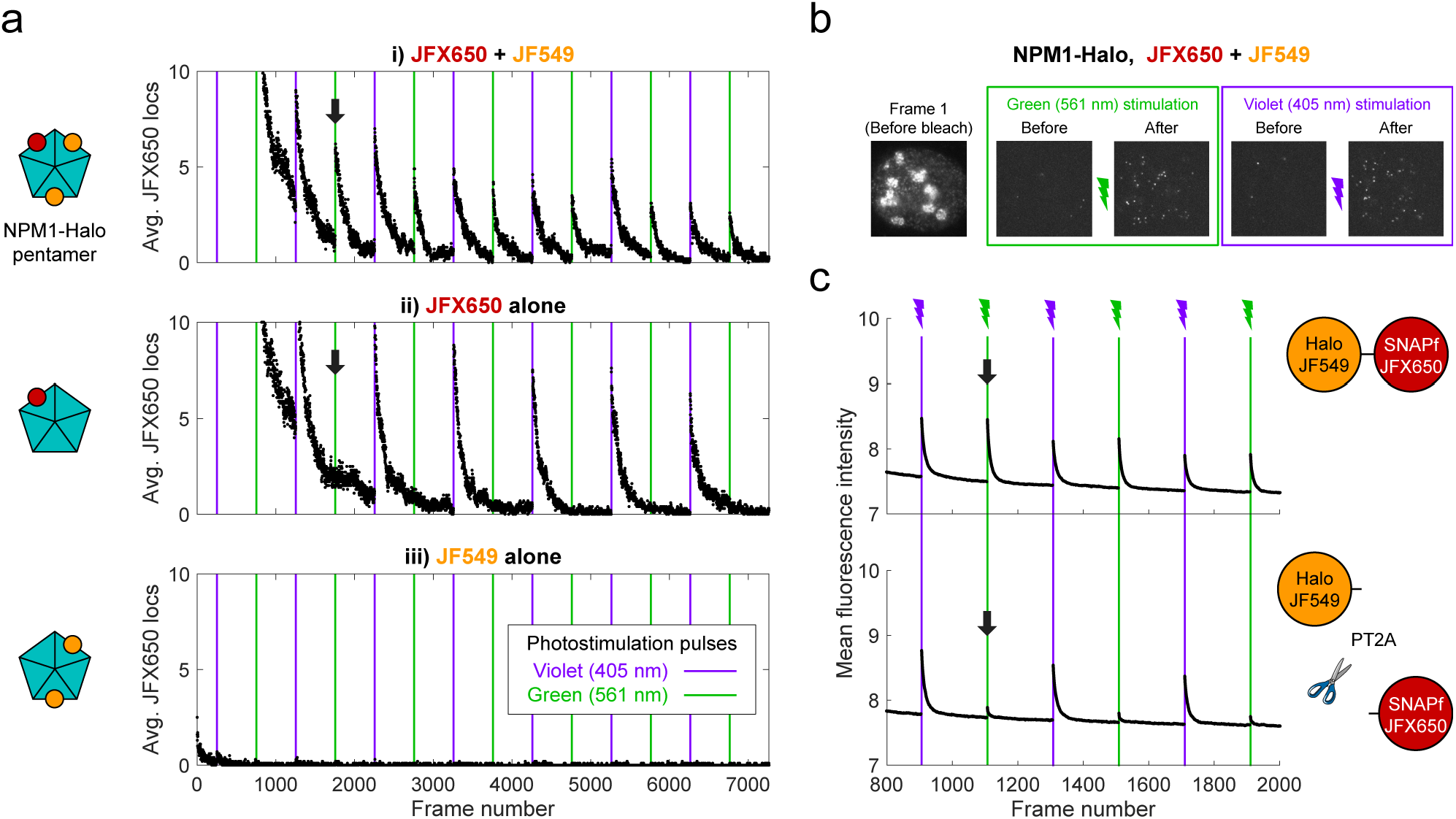
Proximity-assisted photoactivation (PAPA) of JFX650 by JF549. a. Green and violet light reactivate JFX650 through distinct JF549-dependent and JF549-independent mechanisms. Left column: Schematic of NPM1 pentamers in heterozygously tagged NPM1-Halo U2OS cells labeled with JF549 (orange) and/or JFX650 (red). Right column: Average number of localizations in the JFX650 channel as a function of frame number. JFX650 molecules were excited with red (633 nm) light, interspersed with 7-ms pulses of violet (405 nm) and green (561 nm) light (violet and green vertical lines). Reactivation of JFX650 by green light required labeling with JF549 (compare black arrows in (i) and (ii)). b. Sample images of a single cell in the JFX650 channel. Leftmost panel: First movie frame prior to fluorophore bleaching/shelving. Green and violet boxes: Maximum intensity projection of all frames immediately before and after green and violet stimulation pulses, showing reactivation of molecules from the dark state. c. Average fluorescence intensity in the JFX650 channel as a function of frame number in cells expressing a Halo-SNAPf fusion with a flexible linker (top panel; N = 40 cells) or a tandem P2A-T2A self-cleaving peptide between Halo and SNAPf (PT2A; bottom panel; N = 20 cells). Halo was labeled with JF549-HTL and SNAPf with JFX650-STL. Reactivation by violet light pulses (violet lines) occurred in both cases, but reactivation by green light pulses (green lines) was mostly eliminated by the self-cleaving peptide (compare black arrows). Raw intensity traces are displayed without background subtraction.

Because double-labeling of NPM1-Halo pentamers is expected to bring JF549 and JFX650 close together (Fig. 1a(i), right panel), we asked whether proximity of the dyes is required for reactivation. To test this, we expressed fusions of Halo and SNAPf separated by either a short flexible linker (Halo-SNAPf) or a tandem P2A-T2A self-cleaving peptide (Halo-PT2A-SNAPf)^21^ in U2OS cells (Fig. 1c); labeled cells with JF549-HTL and JFX650 SNAP tag ligand (JFX650-STL); and imaged JFX650 with red light interspersed with alternating short pulses of violet and green light. While violet reactivation was similar for both constructs, green reactivation was substantially greater for Halo-SNAPf than for Halo-PT2A-SNAPf, implying that proximity of the two dyes facilitates reactivation by green light (Fig. 1c). Thus, we term this phenomenon **proximity-assisted photoactivation (PAPA)**. We will call the dye that undergoes reactivation the “**receiver**” and the dye whose excitation induces reactivation the “**sender**”. Also, we will adopt the terms “**shelving**” for conversion of the receiver into the dark state^15,22^ and **“direct reactivation” (DR)**for reactivation by violet light^18^.

Conjugating JFX650 to SNAPf instead of Halo led to more efficient shelving in the dark state, as evidenced by a faster decline in fluorescence during red illumination and greater subsequent reactivation by violet light (Supplementary Fig. 1a). Kinetic measurements indicate that about 10% of JFX650-SNAPf molecules enter the dark state under our experimental conditions (Supplementary Fig. 1b). DR by violet light precluded subsequent PAPA by green light, and vice versa, implying that both wavelengths reactivate the same dark state (Supplementary Fig. 1c-d). We tested other fluorophore pairs and found that PAPA occurred when tetramethylrhodamine (TMR), Janelia Fluor X 549 (JFX549), or Janelia Fluor 526 were used as the sender, or when JF646 or JFX646 were used as the receiver (Supplementary Fig. 2).

### Distance-dependence of PAPA

To investigate how PAPA depends on sender-receiver distance, we generated fusion transgenes in which Halo and SNAPf were separated by 0, 1, 3, 5, or 7 repeats of the titin I91 Ig domain^23^. The distance distribution between the two dyes was estimated for each fusion protein by simulating an ensemble of conformations using PyRosetta (Supplementary Fig. 3a-b)^24,25^. U2OS cells were stably transfected with each transgene, and fluorescence-activated cell sorting (FACS) was used to obtain pools of cells with similar low expression levels of each protein (Supplementary Fig. 3c-d; see Supplementary Note 1).

Cells were labeled with a mixture of JF549-HTL and JFX650-STL and imaged as described above with red light interspersed with alternating pulses of violet light to induce DR and green light to induce PAPA. The ratio of the increase in fluorescence intensity in response to green and violet pulses (the “PAPA/DR ratio”) provides a normalized measure of PAPA efficiency, which corrects for cell-to-cell variability in the labeled protein concentration. For sufficiently short reactivation pulses, the PAPA/DR ratio increased linearly with the green pulse duration (with the violet pulse duration held constant), making it possible to measure relative rate constants by linear fitting (Fig. 2a and 2b, left panel). In parallel, fluorescence lifetime imaging (FLIM) was used to measure FRET between JF549 and JFX650 for the same fusion proteins (Fig. 2b, right panel; see Supplementary Note 2).

**Figure 2:**
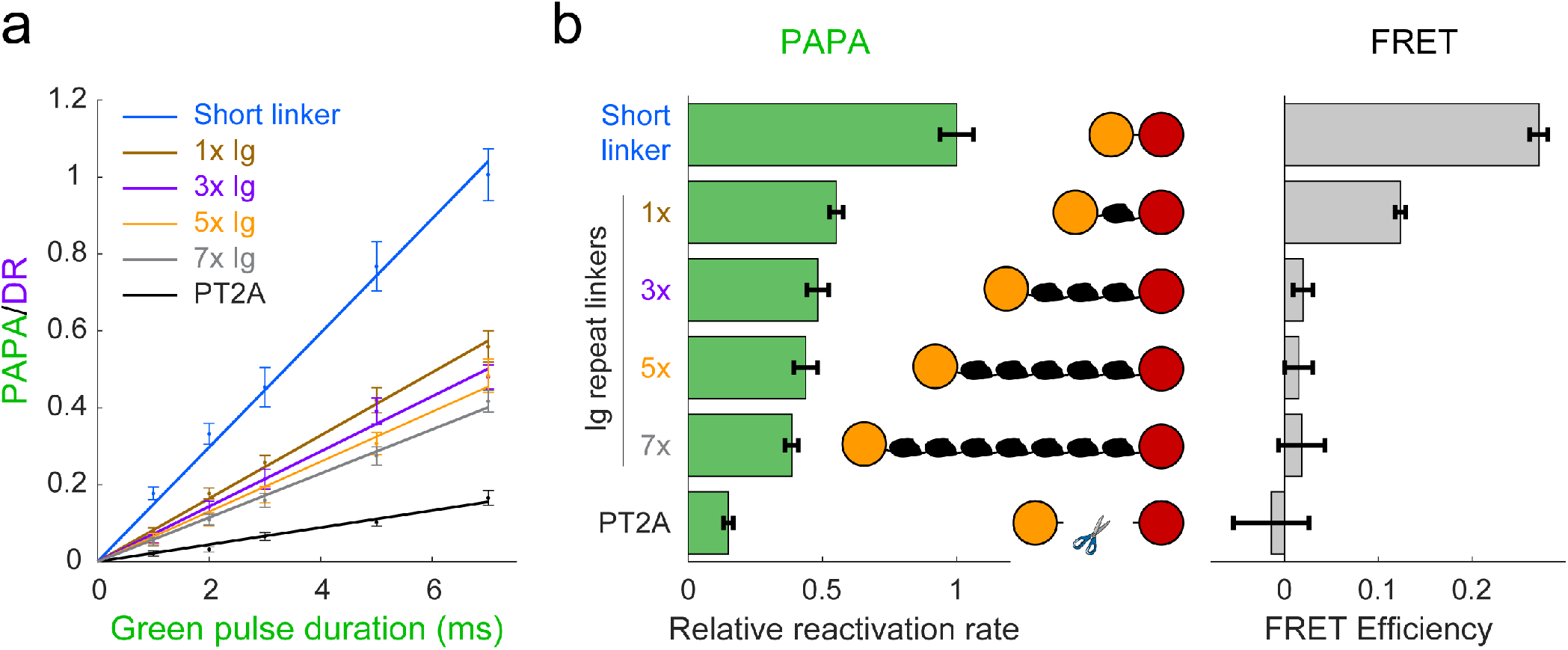
Comparison of distance-dependence of PAPA and FRET. a) PAPA/DR ratio vs. green pulse duration for Halo-SNAPf fusions with a short, flexible linker or linkers containing different numbers of tandem Ig domains. Curves are linear fits (y = ax). Error bars, ± 2 * SE. PT2A, tandem P2A-T2A self-cleaving peptide. b) Left panel: Relative rates of reactivation by PAPA (slope of fits in (a) divided by the slope of the short linker construct). Right panel: FRET efficiency measured using fluorescence lifetime imaging (FLIM).

As predicted by our simulations (Supplementary Fig. 3b), FRET efficiency between JF549 and JFX650 declined sharply with increasing spacer length, from 0.271 ± 0.010 (95% CI) for the short linker to 0.124 ± 0.006 for a single Ig repeat and 0.020 ± 0.010 for three Ig repeats (Fig. 2b, right panel). FRET was essentially undetectable for 5 or 7 Ig repeats and for the PT2A self-cleaving peptide linker (Fig. 2b, right panel). In contrast, PAPA was observed for the 3x, 5x, and 7x Ig linker constructs (Fig. 2a-b). The rate of photoactivation by green light declined gradually with increasing linker length yet was distinguishable from the background rate of the PT2A self-cleaving linker. These results indicate that PAPA has a less stringent dependence on inter-fluorophore distance than FRET.

### Detection of inducible protein-protein interactions using PAPA

Based on the above results, we reasoned that PAPA could be used to detect interaction of two different proteins labeled with SNAPf-JFX650 and Halo-JF549. As a test case, we monitored the rapamycin-inducible interaction of the proteins FRB and FKBP. U2OS cells expressing Halo-FRB and SNAPf-FKBP were labeled with JF549-HTL and JFX650-STL and imaged with alternating green and violet photostimulation as described above (Fig. 3a and Supplementary Fig. 4a). Addition of rapamycin caused a dramatic increase in the ratio of PAPA (green reactivation) to DR (violet reactivation), consistent with ligand-induced dimerization of Halo-FRB and SNAPf-FKBP bringing together JF549 and JFX650 (Fig. 3b-c and Supplementary Fig. 4b).

**Figure 3:**
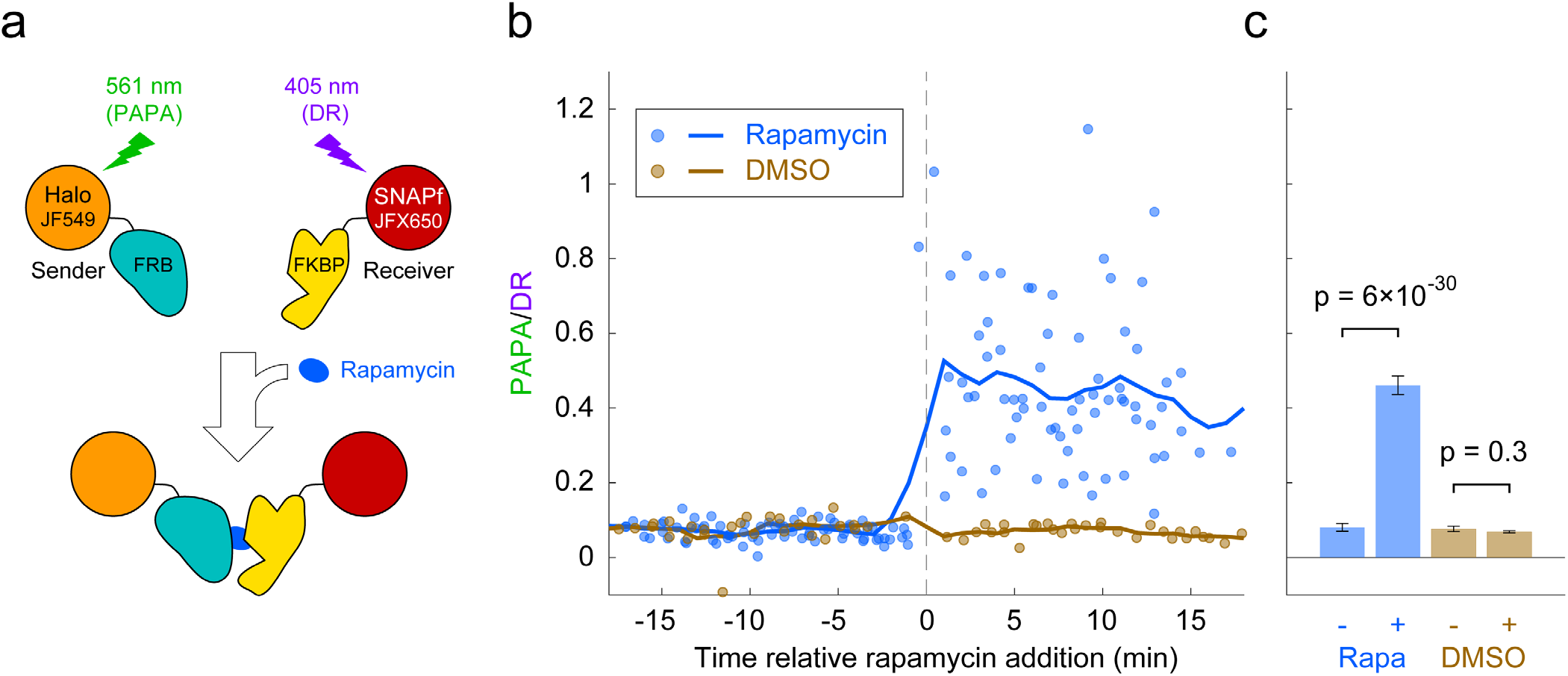
Detection of inducible dimerization using PAPA. a) Halo-FRB was labeled with the sender fluorophore (JF549) and SNAPf-FKBP with the receiver fluorophore (JFX650). After shelving JFX650 with red light, DR and PAPA were alternately induced with pulses of violet and green light, respectively. Midway through the experiment, cells were treated with rapamycin to induce FRB-FKBP dimerization or with dimethylsulfoxide (DMSO) solvent as a negative control. b) Ratio of fluorescence increase due to PAPA (green reactivation) and DR (violet reactivation) as a function of time after rapamycin addition. Blue, rapamycin. Brown, DMSO solvent-only control. Individual data points represent single cells; solid lines show a 2-min moving average. c) Average PAPA/DR ratio before (-) and after (+) addition of rapamycin (Rapa) or DMSO. Total number of cells: 75 before and 74 after rapamycin, 30 before and 30 after DMSO. Error bars, ± 2*SEM. Statistical significance was calculated using a 2-tailed t-test.

### PAPA optically enriches a subset of molecules in defined 2-component mixtures

We next asked whether PAPA can be used to spotlight a sub-population of receiver molecules close to sender molecules. As a simple test case, we analyzed defined mixtures of two proteins—one labeled with JFX650 only, and a second labeled with both JFX650 and JF549—and investigated whether PAPA could optically enrich the double-labeled component to distinguish its properties in single-molecule imaging.

First, we co-expressed SNAPf-tagged histone H2B (SNAPf-H2B), which is predominantly chromatin-bound, along with a Halo-SNAPf fusion with a nuclear localization sequence (Halo-SNAPf-3xNLS), which is mostly unbound (Fig. 4a)^2,26^. Cells were incubated with JFX650-STL and JF549-HTL to label Halo-SNAPf-3xNLS with both JFX650 and JF549 and SNAPf-H2B with JFX650 alone (Fig. 4a, Supplementary Fig. 5a). JFX650 fluorophores were thoroughly photobleached/shelved using a 10-s pulse of intense red light, after which JFX650 was imaged with red light interspersed with pulses of green and violet light. After localizing and tracking single molecules, we separated trajectories occurring after a green pulse (PAPA trajectories) from those occurring after a violet pulse (DR trajectories) and applied a recently developed Bayesian state array SPT (saSPT) algorithm^2^ to infer the underlying distribution of diffusion coefficients for each set of trajectories (its “diffusion spectrum” for short). As predicted, diffusion spectra revealed two peaks, one corresponding to bound molecules (D = 0.01 μm^2^/s, the minimum value in the state array), and one corresponding to freely diffusing molecules (D = 8.3 μm^2^/s). PAPA trajectories were enriched for freely diffusing molecules compared to DR trajectories, as expected if PAPA selectively reactivates JF549/JFX650 double-labeled Halo-SNAPf-3xNLS molecules (Fig. 4b). Next, PAPA and DR trajectories from individual cells were re-analyzed using a 2-state model with bound (D = 0.01 μm^2^/s) and free (D = 8.3 μm^2^/s) states. Consistent with the ensemble analysis, PAPA trajectories had a lower bound fraction than DR trajectories in every cell (Fig. 4c). The same trend is apparent from comparison of particle displacement histograms and raw trajectories (Supplementary Fig. 6a-b).

**Figure 4:**
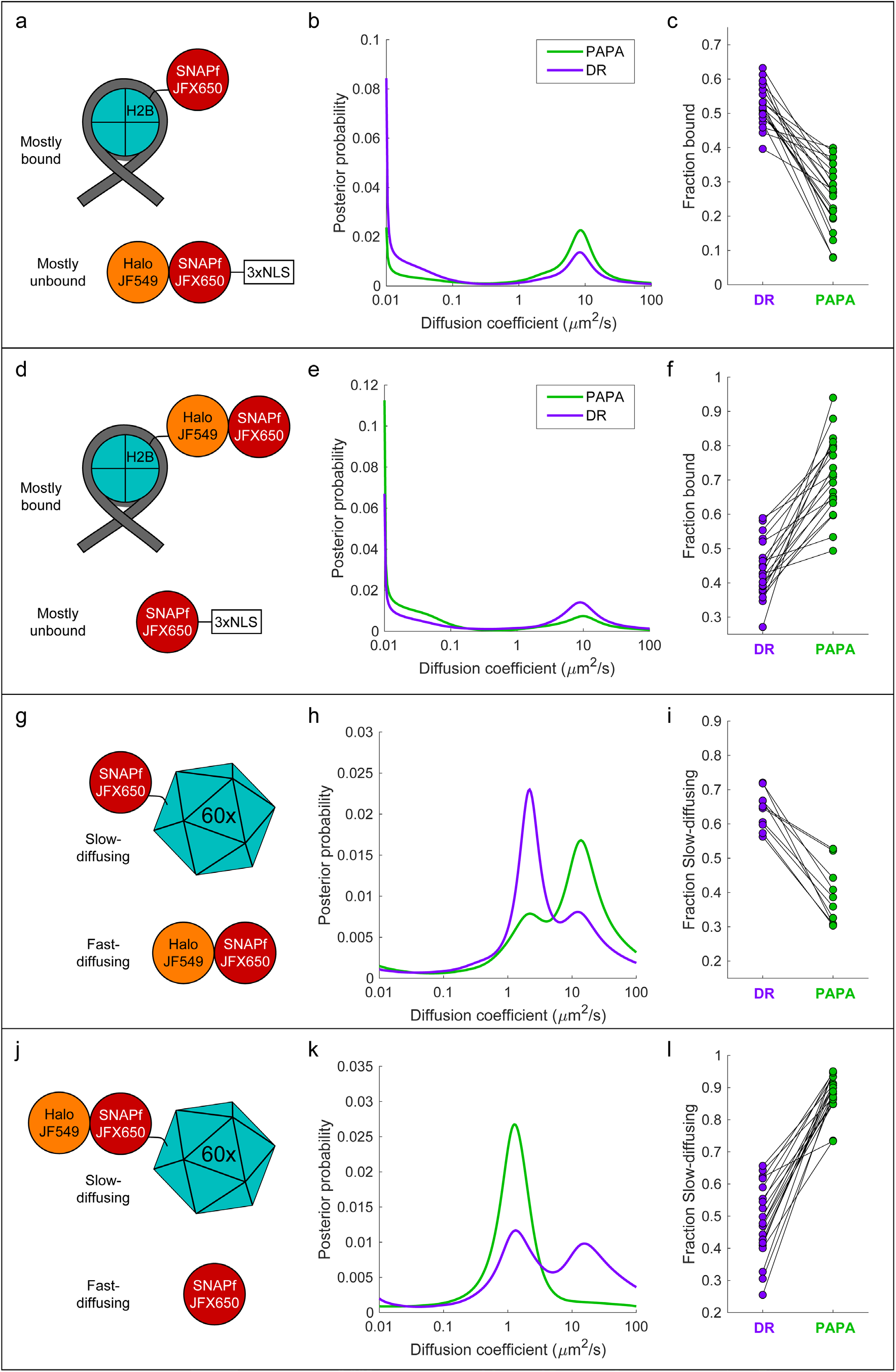
“Unmixing” of defined 2-component mixtures using PAPA. Left column (a,d,g,j): Schematic of different defined mixtures of two labeled proteins, in which one protein is labeled with JFX650 only and the other is labeled with both JFX650 and JF549. In (g) and (j), each subunit of the 60-mer is fused to SNAPf or Halo-SNAPf, though only one label is displayed for clarity. Center column (b,e,h,k): Inferred diffusion spectra of PAPA (green-reactivated) and DR (violet-reactivated) trajectories pooled from 20 cells (b,e,k) or 10 cells (h). Right column: Fraction bound (c,f) or fraction slow-diffusing (i,l) of PAPA and DR trajectories from individual cells, obtained from fits to a 2-state model (c,f) or 3-state model (i,l). Paired, two-tailed t-tests of the comparisons in (c), (f), (i), and (l) showed all differences to be statistically significant with p = 9 × 10^-9^, 8 × 10^-8^, 1 × 10^-5^, and 4 × 10^-11^, respectively.

To exclude the possibility that enrichment of unbound molecules arose from a systematic bias in our method, we repeated the experiment with the reciprocal mixture of Halo-SNAPf-H2B and SNAPf-3xNLS (Fig. 4d). As expected, the opposite trend was observed: PAPA trajectories were enriched in bound molecules compared to DR trajectories, both across an ensemble of cells and at the single-cell level (Fig. 4e-f and Supplementary Fig. 6c-d). As a further control, we analyzed cells expressing JF549-HTL/JFX650-STL double-labeled Halo-SNAPf-3xNLS or Halo-SNAPf-H2B alone. As expected, PAPA and DR trajectories displayed virtually identical diffusion spectra for these individual components (Supplementary Fig. 7a-f).

To test whether PAPA can also distinguish a mixture of diffusing components, we co-expressed fast-diffusing cytosolic Halo-SNAPf with a SNAPf-tagged synthetic protein that forms large, slowly diffusing 60-mers^27^ (Fig. 4g). As expected, diffusion spectra had two peaks corresponding to slow-diffusing (SNAPf-60-mer) and fast-diffusing (Halo-SNAPf) components (Fig. 4h). Compared to DR trajectories (violet curve), PAPA trajectories (green curve) were strongly enriched in the fast-diffusing subpopulation, consistent with selective reactivation of the double-labeled Halo-SNAPf protein by green light (Fig. 4h). The same trend was observed in single-cell reanalysis, displacement histograms, and raw trajectories (Fig. 4i, Supplementary Fig. 6e-f). The enrichment of the fast-diffusing population was not absolute, as a slow-diffusing shoulder peak was still observed among PAPA trajectories (Fig. 4h, green curve; see Discussion). As before, swapping SNAPf and Halo-SNAPf labels yielded the opposite trend, both at the ensemble and single-cell level (Fig. 4j-l and Supplementary Fig. 6g-h). Enrichment of the 60-mer peak by PAPA is especially pronounced in this case (compare green and violet curves in Fig. 4k), which may reflect reactivation of a receiver molecule by any of several neighboring sender molecules within a 60-mer.

Taken together, these results demonstrate that proximity-assisted photoactivation can be used to enrich a subpopulation of molecules in which a receiver fluorophore (e.g., JFX650) is in proximity to a sender fluorophore (e.g., JF549), thereby revealing the distinct properties of this subpopulation at both the ensemble and single-cell level.

### Distinguishing the properties of androgen receptor monomers and dimers in single cells

As a proof-of-concept biological application, we tested whether PAPA could be used to detect ligand-induced self-association of mammalian androgen receptor (AR) and distinguish the properties of AR monomers and dimers/oligomers. First, we stably co-expressed SNAPf and Halo fusions of mouse AR in U2OS cells (which express very little endogenous AR^28^), labeled the two proteins with a mixture of JFX650-STL and JF549-HTL (Supplementary Fig. 8a), and measured the PAPA/DR ratio by quantifying changes in JFX650 fluorescence intensity in response to alternating green and violet stimulation as described above. As expected, treatment with the androgen dihydrotestosterone (DHT) led to an increase in the ratio of PAPA to DR over the course of several minutes (Fig. 5b and Supplementary Fig. 8b), consistent with the two fluorophores being brought together by ligand-induced interaction between SNAPf-mAR and Halo-mAR.

**Figure 5:**
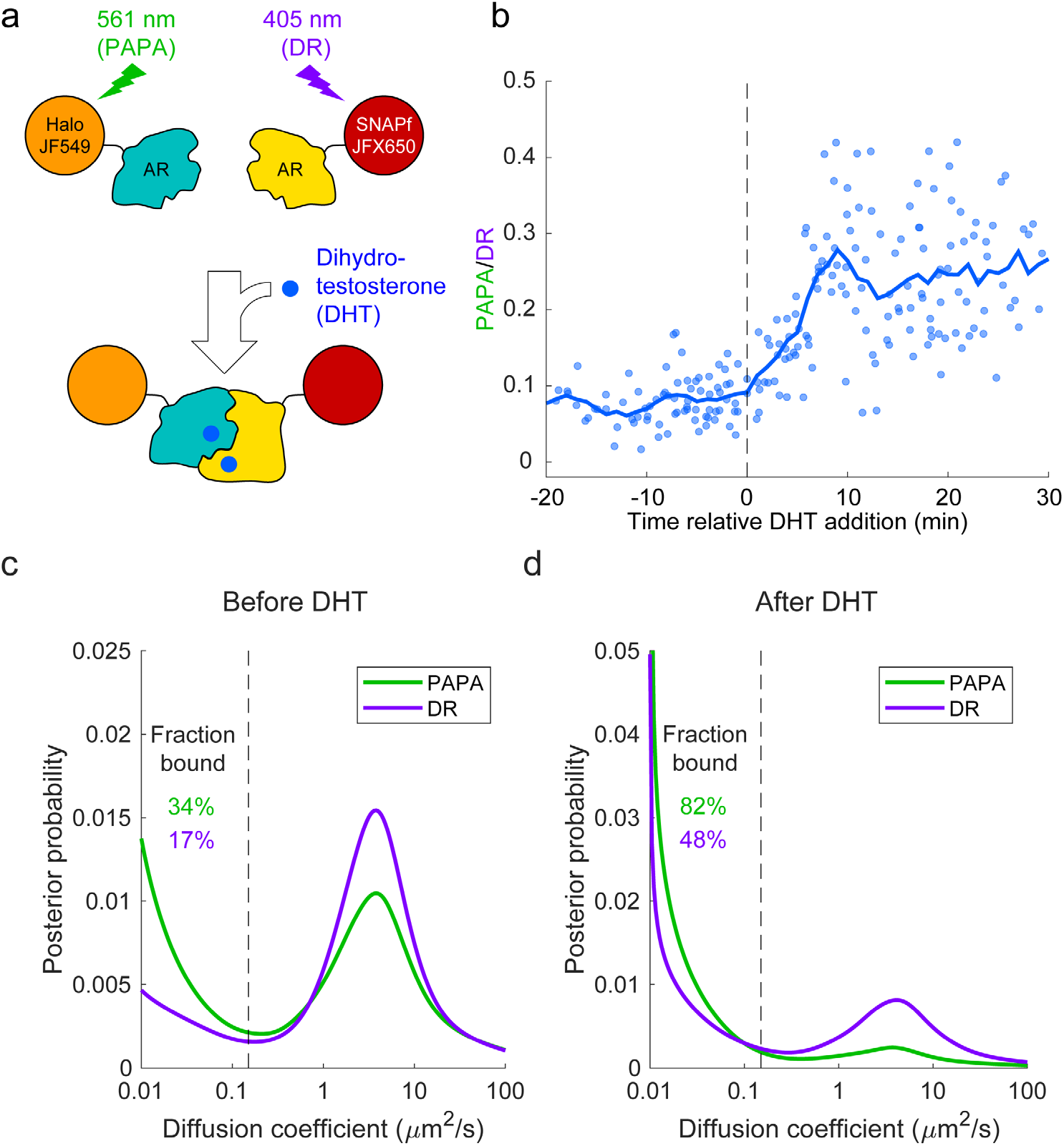
Analysis of mammalian androgen receptor using PAPA-SPT. a) Schematic of DHT-induced dimerization of JF549-Halo-mAR and JFX650-SNAPf-mAR. b) PAPA/DR ratio as a function of time relative DHT addition. c-d) Diffusion spectra of PAPA and DR trajectories. c) Before addition of DHT; N = 55 cells. d) After addition of DHT to a final concentration of 10 nM; N = 81 cells. Fraction bound was quantified by integrating the portion of each curve below D = 0.15 μm^2^/s (vertical dashed line).

Next, we combined PAPA with single-molecule imaging to assess how self-association influences diffusion and chromatin binding by AR. Consistent with previous biochemical and live-cell imaging experiments, addition of DHT caused an increase in the overall bound fraction of AR (Fig. 5c-d)^29–31^. Strikingly, PAPA trajectories had a higher bound fraction than DR trajectories, both before and after addition of DHT (Fig. 5c-d). This is consistent with an increase in the affinity of AR for specific DNA sequence motifs upon self-association. Moreover, PAPA revealed that a subset of AR molecules self-associated and bound chromatin with elevated affinity even prior to addition of exogenous androgen. Thus, PAPA can be applied to monitor regulation of a biologically important protein-protein interaction in live cells, discern its effect on chromatin binding, and reveal the existence of molecular subpopulations.

## Discussion

Here we have described a novel and useful property of rhodamine dyes, proximity-assisted photoactivation (PAPA), in which excitation of a “sender” fluorophore (e.g., JF549) reactivates a nearby “receiver” fluorophore (e.g., JFX650) from a dark state. By enabling targeted reactivation of receiver fluorophores near a sender fluorophore, PAPA provides a new way to detect molecular interactions in live cells (Fig. 6): First, Halo-tagged proteins are labeled with the sender fluorophore and SNAPf-tagged proteins with the receiver fluorophore. Second, cells are illuminated with intense red light to shelve receiver fluorophores in the dark state. Third, alternating pulses of green and violet light are applied to reactivate receiver fluorophores by PAPA and DR, respectively, and these reactivated fluorophores are imaged using red illumination. The ratio of green to violet reactivation provides a measure of protein-protein interaction, while analysis of green-reactivated and violet-reactivated single-particle trajectories makes it possible to compare the overall population of receiver-labeled molecules (violet; DR) to those physically associated with sender-labeled molecules (green; PAPA).

**Figure 6:**
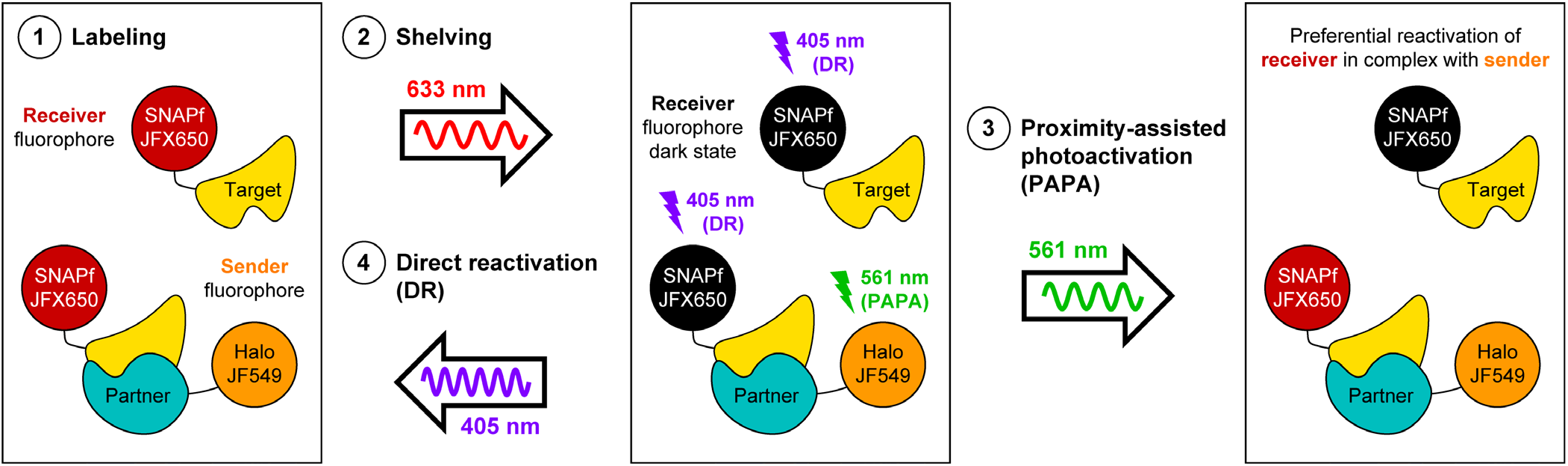
Using PAPA to spotlight protein-protein interactions. 1) Label a SNAPf-tagged Target protein with a receiver fluorophore (e.g., JFX650) and a Halo-tagged Partner protein with a sender fluorophore (e.g., JF549). 2) Shelve the receiver fluorophore in the dark state using intense 633 nm illumination. Image receiver molecules with 633 nm light while alternately illuminating with 3) pulses of 561 nm light to induce proximity-assisted photoactivation (PAPA) of receiver-labeled Target molecules in complex with sender-labeled Partner molecules, and 4) pulses of 405 nm light to induce direct reactivation (DR) of receiver fluorophores, independent of proximity to the sender.

While the physical mechanism underlying PAPA remains unclear, its more flexible distance-dependence than either FRET (Fig. 2) or cyanine dye photoswitching^9^ suggests a distinct process. One hypothesis is that the excited sender reacts with some other molecule in the cell, producing a short-lived chemical species that diffuses a limited distance to react with and reactivate the receiver dark state.

PAPA can complement other techniques for monitoring molecular interactions. Although single-molecule FRET (smFRET) is useful for measuring distances between fluorophores *in vitro*, PAPA provides multiple advantages and opportunities for live-cell imaging: First, PAPA circumvents the tradeoff between labeling density and spectral crosstalk inherent in smFRET. It is impractical to detect molecular interactions in cells by sparsely labeling both the FRET donor and acceptor, as double-labeled complexes will be vanishingly rare. Attempting to solve this problem by densely labeling either the donor or the acceptor creates the new problem of fluorescence bleed-through from the densely labeled channel, which may be orders of magnitude brighter than signal from the sparsely labeled channel. In PAPA, sender and receiver excitation occur at different times, eliminating fluorescence bleed-through from the sender into the receiver channel. Hence, one interacting partner can be sparsely labeled with the receiver and the other densely labeled with the sender, permitting efficient detection of double-labeled complexes. Second, PAPA has a conveniently longer working distance than FRET (Fig. 2), which might be extended further by elongating the linkers between Halo/SNAPf and the protein of interest. Because photoactivation is an all-or-nothing event, a signal can in principle be detected if even a fraction of linker conformations orients the dyes close enough together for PAPA to occur.

Although PAPA significantly enriches for complexes double-labeled with sender and receiver, its selectivity is not perfect. A background level of PAPA was still observed when Halo and SNAPf were separated by a self-cleaving peptide tag (Fig. 1c, 2a-b), which cannot be explained either by incomplete cleavage (Supplementary Fig. 3c-d) or by direct reactivation of JFX650 by 561 nm light (Fig. 1a and Supplementary Fig. 2d). Moreover, some “contamination” of PAPA trajectories with single-labeled molecules was evident in experiments with defined 2-component mixtures (e.g., slower-diffusing peak in green curve of Fig. 4h). This nonspecific background may arise from multiple sources: First, even molecules that are not physically associated come into proximity by chance at some rate. Indeed, when JFX650-labeled cells were bathed in high concentrations of free JF549 dye, reactivation by green light occurred in proportion to the JF549 concentration (Supplementary Fig. 9). Nonspecific background is thus expected to be greater when sender-labeled proteins are expressed at high levels. Second, dark-state fluorophores spontaneously reactivate at a low basal rate even without photostimulation^15,16^. Third, although we chose a time interval between green and violet pulses sufficient to bleach or re-shelve most reactivated fluorophores, it is possible that a small fraction of fluorophores reactivated by a violet pulse survived until the subsequent green pulse. Modeling of these different background contributions will be required to quantify more precisely the characteristics of interacting and non-interacting molecular subpopulations.

Notwithstanding these technical imperfections, PAPA has the potential to open new experimental routes toward understanding the dynamics of protein complexes in live cells. Our results show that PAPA can be used to detect potentially transient protein-protein interactions (Fig. 3 and Supplementary Fig. 4) and to infer the composition of different peaks in single-molecule diffusion spectra (Fig. 4 and Supplementary Fig. 6). A proof-of-concept application to mammalian AR revealed at a single-cell level the relationship between AR self-association and chromatin binding (Fig. 5 and Supplementary Fig. 8). Future applications of PAPA could include measuring differences in the chromatin residence time of different transcriptional subcomplexes, detecting transient interactions mediated by low-complexity domains, or assessing the consequences of posttranslational modifications such as SUMOylation. Although the current study involved labeled proteins, PAPA could potentially be used to detect interactions between other biomolecules as well. By revealing the distinct features of specific molecular complexes, PAPA will provide a powerful new tool to probe biochemical mechanisms in live cells.

## Methods

### Cell culture

U2OS cells were grown in Dulbecco’s modified Eagle’s medium (DMEM) with 4.5 g/L glucose (ThermoFisher # 10566016), 10% fetal bovine serum (FBS) and 100 U/ml penicillin-streptomycin (ThermoFisher #15140122) at 37°C and 5% CO_2_. Phenol red-containing medium was used for propagation of cells, while phenol red-free medium (ThermoFisher # 21063029) was used to minimize fluorescence background in imaging experiments.

### Cloning

Ig linkers were subcloned from a previously described plasmid containing repeats of the titin I91 Ig domain, which were codon-shuffled to prevent recombination^23^. The various Halo and SNAPf fusion constructs described in this paper were generated by PCR and isothermal assembly, and all constructs were completely sequenced before use. Two-component expression plasmids included a codon-shuffled SNAPf-3xNLS-T2A-P2A cassette that was ordered as a gBlock from Integrated DNA Technologies (IDT). All plasmid sequences are available at https://gitlab.com/tgwgraham/papa_paper_plasmids.

### Stable transformation of cells and selection of clonal lines

To generate stable lines by PiggyBac integration, U2OS cells from a confluent 10-cm plate were trypsinized, resuspended in DMEM, and divided between two 15-ml conical tubes. Cells were centrifuged for 2 min at 200 *g*, and the medium was aspirated and replaced with 100 μl of Lonza Kit V transfection reagent (82 μl of Kit V solution and 18 μl of Supplement I; Cat. # VCA-1003) containing 1 μg of the donor plasmid and 1 μg of Super PiggyBac transposase plasmid. The cell suspension was transferred to an electroporation cuvette and electroporated using program X-001 on an Amaxa Nucleofector II (Lonza). Cells in the cuvette were mixed with 300 μl of DMEM, and 100 μl of the cell suspension was diluted in 10 ml of DMEM in a 10-cm plate. After allowing cells to grow for 1-2 days, selection was initiated by adding puromycin to a final concentration of 1 μg/ml.

To generate clonal cell lines of Halo-mAR + SNAPf-mAR and FKBP-SNAPf-3xNLS + FRB-Halo-3xNLS, a polyclonal pool of stably transfected cells from a 10-cm plate was labeled with a mixture of 50 nM JF549 SNAP tag ligand and 50 nM JFX650 Halo tag ligand, and fluorescence-activated cell sorting (FACS) was used to sort single cells expressing both proteins into separate wells of a 96-well plate. For pTG800 (3xFlag-Halo-SNAPf-3xNLS-T2A-P2A-H2B-SNAPf-3xNLS), single-cell clones were obtained by limiting dilution into 96-well plates. Polyclonal pools of stably transfected cells were used for the other 2-component and 1-component experiments in Fig. 4 and Supplementary Fig. 6 and 7.

For the experiments in Fig. 2 and Supplementary Fig. 3, FACS was used to obtain polyclonal pools of U2OS cells expressing a low level of each Halo-linker-SNAPf construct. Confluent 10-cm plates of cells were stained with 50 nM each of JF549-STL and JFX650-HTL, and cells were sorted using the same intensity gate in the JFX650-Halo channel. Cells expressing pTG820 (Halo-3x Ig-SNAPf) and pTG828 (Halo-5x Ig-SNAPf) were sorted on a different day using the intensity of the previously sorted pTG747/U2OS pool to define a gate in the JFX650-Halo channel.

### Visualization of fluorescently labeled proteins by SDS-PAGE

Cells in either 10-cm plates or 6-well plates were labeled with 500 nM of the indicated HTL or STL ligand for 1 h at 37°C, washed twice with 1x PBS, trypsinized, and resuspended in DMEM. Cells were counted using a Countess 3 FL cell counter (Invitrogen), pelleted by centrifugation for 2 min at 200 *g*, and frozen at −80°C. Lysates were prepared by addition of 1 ml (for 10-cm plates) or 200 μl (for 6-well plates) of SDS lysis buffer without dye^32^. Each lysate was passed through a 26-gauge needle 10 times to reduce its viscosity.

Custom 8-well, 1.5 mm combs for SDS-PAGE were 3D-printed using an AnyCubic Photon 3D printer (model files available at https://gitlab.com/tgwgraham/gel-combs). For the gels in Supplementary Fig. 3c-d, samples of cell lysate corresponding to 100,000 cells were separated on a 10% SDS-PAGE gel, which was imaged on a Pharos FX imager (BioRad) using the “low-intensity” setting in the Cy3 channel. Cell lysate corresponding to 60,000 cells was loaded per lane of the gels in Supplementary Fig. 4, 5, and 8. The gel in Supplementary Fig. 5a was imaged on a Pharos FX imager (BioRad) using the “low-intensity” setting in the Cy5 channel. The gels in Supplementary Fig. 4a, 5b, and 8a were imaged in the 700 nm channel on an Odyssey imager (LI-COR) at 169 μm resolution with the “medium” quality setting and a z-offset of 0.5 mm. Precision Plus Protein All Blue Prestained Protein Standards (BioRad #1610373) were used as molecular weight standards for all gels.

### Live-cell single-molecule imaging

One day prior to imaging, 25-mm No. 1.5H glass coverslips (Marienfeld, #0117650) were immersed in isopropanol, transferred with forceps to 6-well plates, and aspirated thoroughly to remove all traces of isopropanol. Cells were trypsinized, counted using a Countess 3 FL cell counter (Invitrogen), centrifuged for 2 min at 200 *g*, resuspended in phenol red-free DMEM, and plated at a density of 5 × 10^5^ cells per well. Just prior to imaging, cells were incubated with Janelia Fluor HaloTag and SNAP tag ligands in phenol red-free DMEM for 15 min at 37°C, washed twice with 1x phosphate buffered saline, and destained for at least 15 min in phenol red-free DMEM. The following dye concentrations were used for staining:

- 10 nM JF549-HTL and/or 250 pM JFX650-HTL for Fig. 1a-b.
- 50 nM JF549-HTL and 5 nM JFX650-STL for Fig. 1c, 2a-b, 3, 4, 5, and Supplementary Fig. 1b-d, 4b, 6, 7, 8, and 9 (dashed blue line).
- 50 pM JFX650 HTL or 5 nM JFX650-STL for Supplementary Fig. 1a.
- 50 nM JF526/JF549/JFX549/TMR-HTL and/or 5 nM JF646/JFX650-STL for Supplementary Fig. 2.

Coverslips were mounted in a stainless steel Attofluor^™^ Cell Chamber (ThermoFisher #A7816) and covered with 1 ml of phenol red-free DMEM with 10% FBS and penicillin/streptomycin. Cells were imaged using HILO illumination on the microscope described in detail in Ref. ^1^. Laser power densities used for imaging were approximately 52 W/cm^2^ for 405 nm (violet), 100 W/cm^2^ for 561 nm (green), and 2.3 kW/cm^2^ for 633 nm (red). Fluorescence emission was filtered through a Semrock 676/37 bandpass filter.

Cells were imaged at a rate of 7.48 ms/frame. Different experiments employed variations of an illumination sequence with alternating pulses of 633 nm red (R), 561 nm green (G), and 405 nm violet (V) light synchronized to the camera. We use these abbreviations below and indicate the duration of the light pulse in brackets. For instance, “250 R [2 ms]” denotes 250 frames with a 2-ms pulse of red 633 nm illumination per frame. Red illumination was restricted to one 2-ms stroboscopic pulse per frame in single-molecule tracking to reduce the motion blur of moving molecules^26^. The green and violet pulse durations were adjusted in different experiments to maintain a trackable density of localizations after each pulse.

- Fig. 1a-b, 3, 5b, Supplementary Fig. 4b, 8b: 10 cycles of 250 R [2 ms], 1 V [7 ms], 500 R [2 ms], 1 G [7 ms], 250 R [2 ms]. The number of cycles in Fig. 3 and 5b was reduced to 4 and 5, respectively.
- Fig. 1c and Fig. 2: 5 cycles of 100 R [7 ms], 1 V [7 ms] + R [7 ms], 200 R [7 ms], 1 G [7 ms] + R [7 ms], 100 R [7 ms]
- Fig. 4a-f, 5c-d, 8c, and Supplementary Fig. 4, 6, 7a-f: Cells were first illuminated 10 s with 633 nm light to either photobleach or shelve most JFX650 fluorophores and then imaged with 10 cycles of 250 R [2 ms], 1 V [0.5 ms] + R [2 ms], 500 R [2 ms], 1 G [X ms] + R [2 ms], 250 R [2 ms]. Owing to differences in the PAPA efficiency and protein concentration between samples, the green pulse duration (X) was adjusted empirically to obtain a trackable number of localizations following photostimulation. It was set to 0.5 ms for Fig. 4g-l and Supplementary Fig. 6e-h, 7g-I; 2 ms for Fig. 4a-f, 5d, 8c (after DHT), and Supplementary Fig. 6a-d, 7a-f; and 7 ms for Fig. 5c and 8c (before DHT). Because DHT addition greatly increased the PAPA efficiency in AR experiments (Fig. 5b), the green pulse duration was shortened from 7 ms before DHT to 2 ms after DHT in Fig. 5c-d and 8c to keep the number of localizations per frame roughly equivalent and prevent PAPA trajectories from becoming too dense for accurate tracking.
- Supplementary Fig. 1a: 10 cycles of 100 R [7 ms], 1 V [7 ms] + R [7 ms], 100 R [7 ms]. Only the first two cycles are shown in the figure.

### Analysis of ensemble PAPA experiments

For ensemble PAPA experiments (Fig. 1c, 2a-b, 3b-c, 5b and Supplementary Fig. 1, 2, 4b, 8b, 9), custom MATLAB code was used to sum the total intensity of all pixels in the field of view at each frame. Frame-by-frame intensity across multiple movies was averaged to obtain “sawtooth” plots of intensity vs. frame number (Fig. 1c and Supplementary Fig. 1a,c,d, 2, 4b, 8b). The PAPA/DR ratio was calculated for Fig. 2a-b, 3b-c, 5b and Supplementary Fig. 9 by dividing the mean increase in fluorescence intensity induced by green and violet pulses. For Fig. 2a-b, which used an illumination sequence with fewer frames per cycle, the initial green and violet pulse were omitted from the averages to avoid the transient photobleaching/shelving phase at the beginning of each movie.

### Quantification of reactivation kinetics

To measure direct reactivation of JFX650-SNAPf as a function of 405 nm illumination time (Supplementary Fig. 1b), cells expressing Halo-SNAPf-3xNLS (pTG747) were stained with 5 nM JFX650 STL and 50 nM JF549 HTL and imaged at 7.48 ms/frame using a 5-phase protocol: 1) 20 frames with 1-ms pulses of 633 nm (intensity measurement before bleaching/shelving), 2) 400 frames with 7-ms pulses of 633 nm (bleaching/shelving), 3) 20 frames with 1-ms pulses of 633 nm (intensity measurement after bleaching/shelving), 4) N frames with 7-ms pulses of 405 nm (reactivation), 5) 20 frames with 1-ms pulses of 633 nm (intensity measurement after reactivation). The total pixel intensity was summed for all 20 frames in phases (1), (3), and (5), and the fractional reactivation was calculated by subtracting the increase in signal between (3) and (5) by the initial drop in signal between (1) and (3). The number of violet frames in phase 4, N, was varied as indicated by the values on the horizontal axis of Supplementary Fig. 1b. The data were fitted to a single-exponential model.

### FLIM-FRET

Fluorescence lifetime was measured on a Zeiss LSM 980 confocal microscope equipped with a Becker-Hickl SPC-150NX TCSPC module. Cells were labeled with a mixture of 50 nM JFX650-HTL and 50 nM JF549-STL for 1 h at 37°C, or with 50 nM JF549-STL alone as a no-FRET control. After briefly washing twice with 1x PBS, cells were destained for at least 15 min prior to imaging. JF549 was excited using a 562 nm laser, and fluorescence emission was filtered through a Semrock 593/40 bandpass filter. Signal was acquired for 10 s over a 256 x 256 px region centered on each cell nucleus. Raw data were imported into SPCImage (Becker & Hickl) and fitted to a single exponential model without binning pixels. Decay constants and total fluorescence intensities for each pixel were exported in csv format. Custom MATLAB code was used to define nuclear masks by intensity thresholding and determine the mean fluorescence lifetime within the nucleus. FRET efficiency was calculated using the formula E_FRET_ = 1-τ/τ_0_, where τ is the fluorescence lifetime of the sample and τ_0_ is the fluorescence lifetime of cells stained with JF549-STL donor only.

### Measuring mutual “occlusion” of PAPA and DR

To measure whether PAPA precludes DR and vice versa (Supplementary Fig. 1c-d), we first labeled cells expressing Halo-SNAPf-3xNLS (pTG747) with 5 nM JFX650 STL and 50 nM JF549. We then imaged at 7.48 ms/frame in 3 phases: 1) 500 frames of 633 nm light (7 ms pulses), alternating with unrecorded frames with either no illumination (black curves) or 7 ms pulses of 405 nm (violet curves) or 561 nm (green curves) illumination. 2) 20 frames with 7 ms pulses of 405 nm light (Supplementary Fig. 1c) or 561 nm light (Supplementary Fig. 1d). 3) 100 frames of 633 nm light (7 ms pulses). Fluorescence intensity traces were prepared as described in “Analysis of ensemble PAPA experiments” above.

### Analysis of SPT data

Particles were localized and tracked using the quot package (https://github.com/alecheckert/quot) with default settings. Custom MATLAB code (https://gitlab.com/tgwgraham/papacode_v1) was used to extract all trajectory segments occurring within the first 30 frames after pulses of 405 nm light (DR trajectories) and 561 nm light (PAPA trajectories). PAPA and DR trajectories were then separately analyzed using a Bayesian “fixed-state sampler” algorithm (https://github.com/alecheckert/spagl)^2^, which estimates the posterior probability distribution over a fixed array of diffusion coefficients. To allow a side-by-side comparison of the distributions for PAPA and DR trajectories, the same number of trajectories were included in the analysis for each. To this end, trajectories were randomly subsampled without replacement from whichever condition, PAPA or DR, had more trajectories. The fraction bound was calculated in Fig. 4c,f and Supplementary Fig. 7c,f by reanalyzing the data using a reduced 2-state model with diffusion coefficients 0.01 and 8.3 μm^2^/s. The fraction slow-diffusing was calculated in Fig. 4i,l and Supplementary Fig. 7i by fitting to a 3-state model with diffusion coefficients 0.01, 2.1, and 13.2 μm^2^/s (Fig. 4i) or 0.01, 1.3, and 15.8 μm^2^/s (Fig. 4l and Supplementary Fig. 7i). Fraction bound for androgen receptor (Supplementary Fig. 8c) was calculated by fitting to a 2-state model with diffusion coefficients 0.01 and 4.4 μm^2^/s. Diffusion coefficients used in the reduced models correspond to the local maxima of the ensemble distributions (Fig. 4b,e,h,k and 5c-d). Displacement histograms in Supplementary Fig. 6b,d,f,h were tabulated using custom code in MATLAB.

A more streamlined and user-friendly Python module for PAPA-SPT analysis will be maintained at https://gitlab.com/tgwgraham/papacode_v2.

### PyRosetta simulations

Inter-fluorophore distances were computed from simulated structural ensembles of each linker construct generated using PyRosetta. Crystal structures of Halo (PDB: 6u32), SNAPf (PDB: 6y8p), and titin Ig (PDB: 1tit) were used to model structured regions. Regions lacking density were filled in using RosettaRemodel, and co-crystalized fluorophores bound to Halo and SNAPf were used to estimate inter-fluorophore distance^33^. After filling in missing residues, each structure was minimized using the FastRelax protocol, and starting structures were generated by concatenating structured regions using linkers corresponding to those used in experimental constructs. Ensembles were generated using an adapted version of the FastFloppyTail method used for sampling disordered protein regions, in which only residues comprising the inter-domain linkers were allowed to move^25^. The adapted version of the FastFloppyTail algorithm features the addition of the BackrubMover, to allow for motion within a large loop present in SNAPf and facilitate more complete sampling^34^. After application of the adapted FastFloppyTail protocol, resultant structures were minimized using FastRelax. Each ensemble consisted of 100 structures from which inter-fluorophore distances were computed. Scripts used to generate these ensembles along with the input structures and resultant ensembles can be found at https://github.com/jferrie3/FusionProteinEnsemble.

## Acknowledgements

Thanks to Luke Lavis, Samantha Rider, Alec Heckert, John Lis, Philip Versluis, Joe Loparo, Max Staller, and the members of the Tjian-Darzacq group for helpful discussions; to Matt Akamatsu for sharing a plasmid encoding the synthetic 60-mer protein; to Vinson Fan for help with androgen receptor cloning; to the UC Berkeley Flow Cytometry Facility for assistance generating clonal cell lines; and to Holly Aaron (UC Berkeley Molecular Imaging Center) and Ana Robles for assistance with microscopy. T.G. was supported by a postdoctoral fellowship from the Jane Coffin Childs Memorial Fund for Medical Research, J.F. is a Howard Hughes Medical Institute Awardee of the Life Sciences Research Foundation, and R.T. is an investigator of the Howard Hughes Medical Institute.

## Competing interests

R.T. and X.D. are co-founders of Eikon Therapeutics, Inc.

## Notes

https://gitlab.com/tgwgraham/papa_paper_plasmids

https://gitlab.com/tgwgraham/papacode_v1

https://gitlab.com/tgwgraham/papacode_v2

https://gitlab.com/tgwgraham/gel-combs

https://github.com/jferrie3/FusionProteinEnsemble

## References

1. Hansen, A. S. et al. Robust model-based analysis of single-particle tracking experiments with Spot-On. Elife 7, (2018).

2. Heckert, A., Dahal, L., Tjian, R. & Darzacq, X. Recovering mixtures of fast diffusing states from short single particle trajectories. bioRxiv (2021).

3. Chen, Y. et al. Mechanisms Governing Target Search and Binding Dynamics of Hypoxia-Inducible Factors. bioRxiv (2021).

4. Nguyen, V. Q. et al. Spatiotemporal coordination of transcription preinitiation complex assembly in live cells. Mol. Cell 81, 3560–3575.e6 (2021).

5. Quast, R. B. & Margeat, E. Single-molecule FRET on its way to structural biology in live cells. Nat. Methods 18, 344–345 (2021).

6. Hansen, A. S., Pustova, I., Cattoglio, C., Tjian, R. & Darzacq, X. CTCF and cohesin regulate chromatin loop stability with distinct dynamics. Elife 6, (2017).

7. Hansen, A. S., Amitai, A., Cattoglio, C., Tjian, R. & Darzacq, X. Guided nuclear exploration increases CTCF target search efficiency. Nat. Chem. Biol. 16, 257–266 (2020).

8. McSwiggen, D. T. et al. Evidence for DNA-mediated nuclear compartmentalization distinct from phase separation. Elife 8, (2019).

9. Bates, M., Blosser, T. R. & Zhuang, X. Short-range spectroscopic ruler based on a single-molecule optical switch. Phys. Rev. Lett. 94, 108101 (2005).

10. Rust, M. J., Bates, M. & Zhuang, X. Sub-diffraction-limit imaging by stochastic optical reconstruction microscopy (STORM). Nat. Methods 3, 793–5 (2006).

11. Geertsema, H. J. et al. Single-molecule imaging at high fluorophore concentrations by local activation of dye. Biophys. J. 108, 949–956 (2015).

12. Chen, Y., Gu, M., Gunning, P. W. & Russell, S. M. Dense small molecule labeling enables activator-dependent STORM by proximity mapping. Histochem. Cell Biol. 146, 255–66 (2016).

13. van de Linde, S. et al. Photoinduced formation of reversible dye radicals and their impact on super-resolution imaging. Photochem. Photobiol. Sci. 10, 499–506 (2011).

14. Heilemann, M. et al. Subdiffraction-resolution fluorescence imaging with conventional fluorescent probes. Angew. Chem. Int. Ed. Engl. 47, 6172–6 (2008).

15. Grimm, J. B. et al. A general method to improve fluorophores for live-cell and single-molecule microscopy. Nat. Methods 12, 244–50, 3 p following 250 (2015).

16. Tang, X. et al. Kinetic principles underlying pioneer function of GAGA transcription factor in live cells. bioRxiv (2021).

17. Vaughan, J. C., Jia, S. & Zhuang, X. Ultrabright photoactivatable fluorophores created by reductive caging. Nat. Methods 9, 1181–4 (2012).

18. Dempsey, G. T. et al. Photoswitching mechanism of cyanine dyes. J. Am. Chem. Soc. 131, 18192–3 (2009).

19. Gidi, Y. et al. Unifying Mechanism for Thiol-Induced Photoswitching and Photostability of Cyanine Dyes. J. Am. Chem. Soc. 142, 12681–12689 (2020).

20. Grimm, J. B. et al. A General Method to Improve Fluorophores Using Deuterated Auxochromes. JACS Au 1, 690–696 (2021).

21. Liu, Z. et al. Systematic comparison of 2A peptides for cloning multi-genes in a polycistronic vector. Sci. Rep. 7, 2193 (2017).

22. Bretschneider, S., Eggeling, C. & Hell, S. W. Breaking the diffraction barrier in fluorescence microscopy by optical shelving. Phys. Rev. Lett. 98, 218103 (2007).

23. Scholl, Z. N., Josephs, E. A. & Marszalek, P. E. Modular, Nondegenerate Polyprotein Scaffolds for Atomic Force Spectroscopy. Biomacromolecules 17, 2502–5 (2016).

24. Chaudhury, S., Lyskov, S. & Gray, J. J. PyRosetta: a script-based interface for implementing molecular modeling algorithms using Rosetta. Bioinformatics 26, 689–691 (2010).

25. Ferrie, J. J. & Petersson, E. J. A Unified De Novo Approach for Predicting the Structures of Ordered and Disordered Proteins. J. Phys. Chem. B 124, 5538–5548 (2020).

26. Hansen, A. S. et al. Robust model-based analysis of single-particle tracking experiments with Spot-On. Elife 7, (2018).

27. Hsia, Y. et al. Design of a hyperstable 60-subunit protein dodecahedron. [corrected]. Nature 535, 136–9 (2016).

28. Dellal, H. et al. High Content Screening Using New U2OS Reporter Cell Models Identifies Harmol Hydrochloride as a Selective and Competitive Antagonist of the Androgen Receptor. Cells 9, (2020).

29. Schaufele, F. et al. The structural basis of androgen receptor activation: intramolecular and intermolecular amino-carboxy interactions. Proc. Natl. Acad. Sci. U. S. A. 102, 9802–7 (2005).

30. van Royen, M. E., van Cappellen, W. A., de Vos, C., Houtsmuller, A. B. & Trapman, J. Stepwise androgen receptor dimerization. J. Cell Sci. 125, 1970–9 (2012).

31. van Royen, M. E. et al. Compartmentalization of androgen receptor protein-protein interactions in living cells. J. Cell Biol. 177, 63–72 (2007).

32. Cattoglio, C. et al. Determining cellular CTCF and cohesin abundances to constrain 3D genome models. Elife 8, (2019).

33. Huang, P.-S. et al. RosettaRemodel: A Generalized Framework for Flexible Backbone Protein Design. PLoS One 6, e24109 (2011).

34. Smith, C. A. & Kortemme, T. Backrub-Like Backbone Simulation Recapitulates Natural Protein Conformational Variability and Improves Mutant Side-Chain Prediction. J. Mol. Biol. 380, 742–756 (2008).

