## Supplementary Material for "Proximity-assisted photoactivation (PAPA): Detecting molecular interactions in live-cell single-molecule imaging"

### Supplementary Figures

#### Supplementary Fig. 1

**Properties of JFX650 reactivation.**

a) Shelving/bleaching and reactivation of JFX650 bound to Halo (cyan) and SNAPf (black). Relative fluorescence intensity is plotted on the y-axis, averaged over multiple cells (N = 10 for Halo, N = 14 for SNAPf), and frame number is plotted on the x-axis. The frame rate was 7.48 ms/frame. Fluorescence intensity (with red illumination) declined more rapidly for SNAPf-JFX650 than for Halo-JFX650. Violet pulses of 7 ms at frames 101 and 302 induced greater direct reactivation of JFX650-SNAPf than JFX650-Halo.

b) Reactivation of SNAPf-JFX650 as a function of violet pulse duration. SNAPf-JFX650 intensity was measured three times using 20 frames of 1 ms stroboscopic red illumination: 1) before bleaching/shelving, 2) after bleaching/shelving with 400 frames of non-stroboscopic (7 ms/frame) red illumination, and 3) after reactivation by exposure to violet pulses of varying duration. Percent reactivation was calculated by dividing the increase in intensity due to reactivation by the decrease in intensity due to bleaching/shelving. Solid black curve shows a fit to a single-exponential model.

c-d) Mutual occlusion of green and violet reactivation. c) Halo-SNAPf-expressing U2OS cells labeled with JFX650-STL and JF549-HTL were imaged with red illumination alternating with unrecorded frames with either no illumination (black curve) or violet/green illumination (violet/green curves). Integrated intensity in the JFX650 channel is plotted on the y-axis and frame number on the x-axis. JFX650 intensity initially declined for all conditions due to bleaching and shelving. A 20-frame (140-ms) pulse of green light was applied after frame 500 (green rectangle). This reactivated JFX650 that had been exposed to red light only but failed to reactivate JFX650 that had been exposed to alternating red and green or red and violet light. d) Same as (c), but with a 20-frame (140-ms) violet pulse after frame 500. Violet reactivation was strongly reduced by preceding green or violet light exposure. Some violet reactivation was still observed for cells exposed to alternating red and green light (green curve). Although this might indicate the presence of additional dark state(s) that can be reactivated by 405 nm light but not PAPA, it might also reflect incomplete labeling of Halo by JF549 or photobleaching of JF549 by green light.

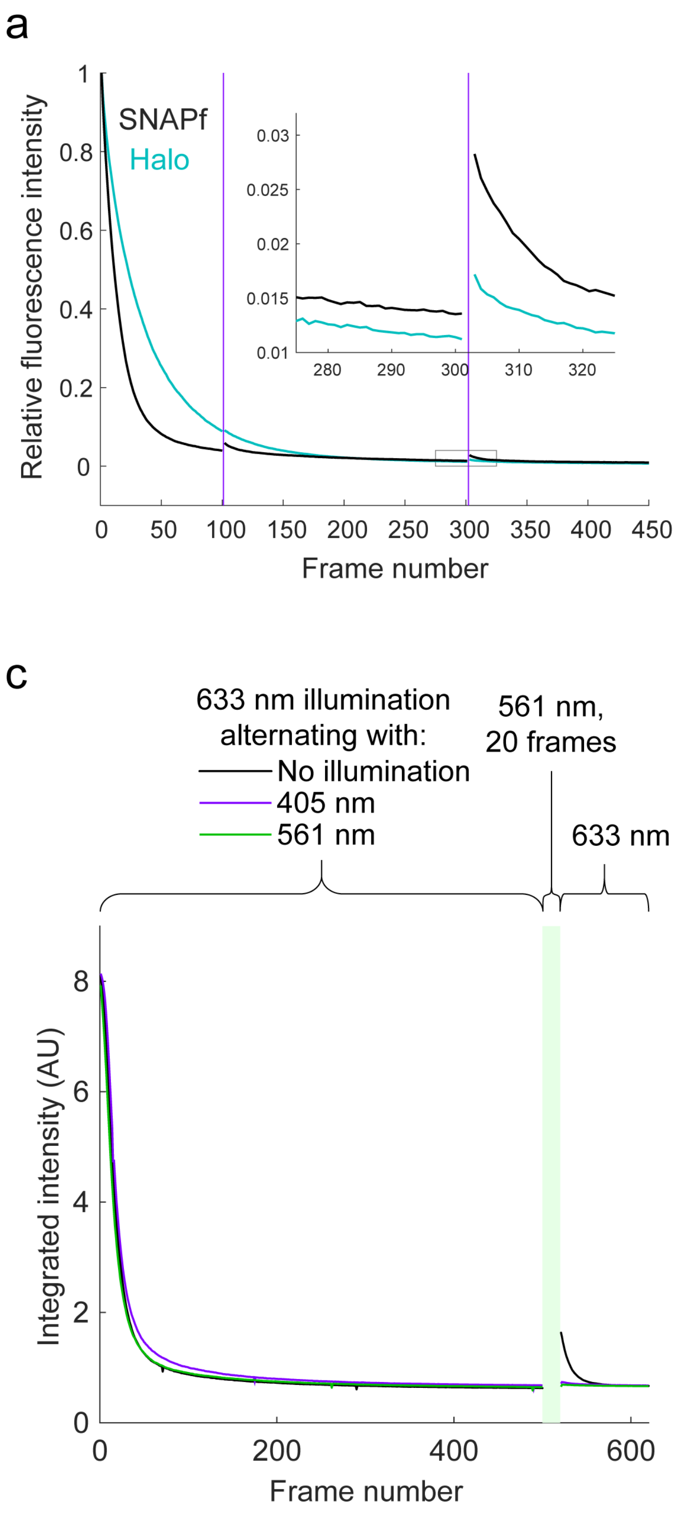

a
SNAPf
Halo
Relative fluorescence intensity
Frame number
b
Fraction reactivated
405 nm pulse duration (s)
Data
Single-exponential fit
$y = a(1 - e^{-kt})$
a = 0.103
k = 19/s
c
633 nm illumination alternating with:
No illumination
405 nm
561 nm
561 nm, 20 frames
633 nm
Integrated intensity (AU)
Frame number
d
633 nm illumination alternating with:
No illumination
405 nm
561 nm
405 nm, 20 frames
633 nm
Integrated intensity (AU)
Frame number

### Supplementary Fig. 2

**PAPA between other sender-receiver pairs.** U2OS cells expressing Halo-SNAPf-3xNLS were labeled with different SNAP tag ligand (STL) and HaloTag ligand (HTL) fluorophore combinations and imaged with red light alternating with 7-ms pulses of green and violet light. Fluorescence intensity averaged over multiple cells is plotted on the vertical axis and frame number is plotted on the horizontal axis. a) Tetramethylrhodamine (TMR)-HTL and JFX650-STL. b) Janelia Fluor X549 (JFX549)-HTL and Janelia Fluor 646 (JF646)-STL. c) Janelia Fluor 526 (JF526)-HTL and JFX650-STL. d) JFX650-STL-only control. e) JF549-only control.

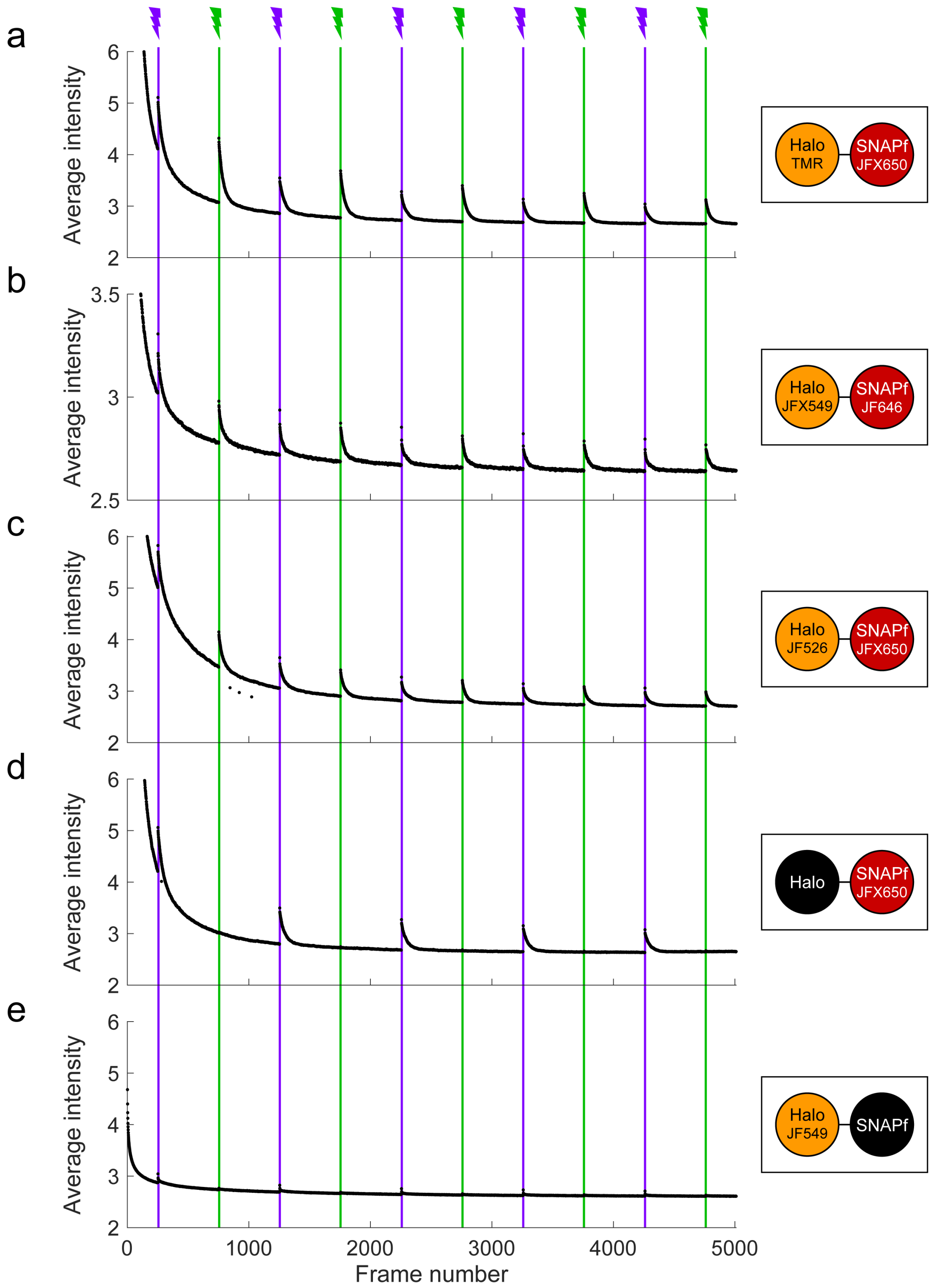

### Supplementary Fig. 3

**Simulations and SDS-PAGE analysis of linker constructs.** a) Four example structures of the Halo-3xIg-SNAPf fusion protein obtained from PyRosetta simulations. Inter-dye distances are indicated for each structure. b) Cumulative distributions of predicted inter-dye distances from Rosetta simulations, with observed (obs.) and predicted (pred.) FRET values. Predicted FRET efficiency was calculated by averaging $E_{FRET} = 1/(1+(R/R_0)^6)$ over all conformations in the ensemble, assuming a theoretical Förster radius of $R_0$ = 58 Å for the JF549-JFX650 pair. c-d) SDS-PAGE analysis of linker constructs expressed in U2OS cells and labeled with JF549-HTL (c) or JF549-STL (d). The amount of cell lysate loaded in each lane corresponds to 100,000 cells. Lookup table is set between 0 and 3000 counts for (c) and the top image in (d). The bottom image in (d) is the same gel with lookup table set between 0 and 800 counts to highlight faint bands. MW, molecular weight markers in kilodaltons. *, nonspecific bands. ◊, unbound dye. A ladder of smaller fragments is present below the full-length protein for the larger linkers; these are predominantly labeled by Halo ligand but not SNAP ligand (see Supplemental Note 1).

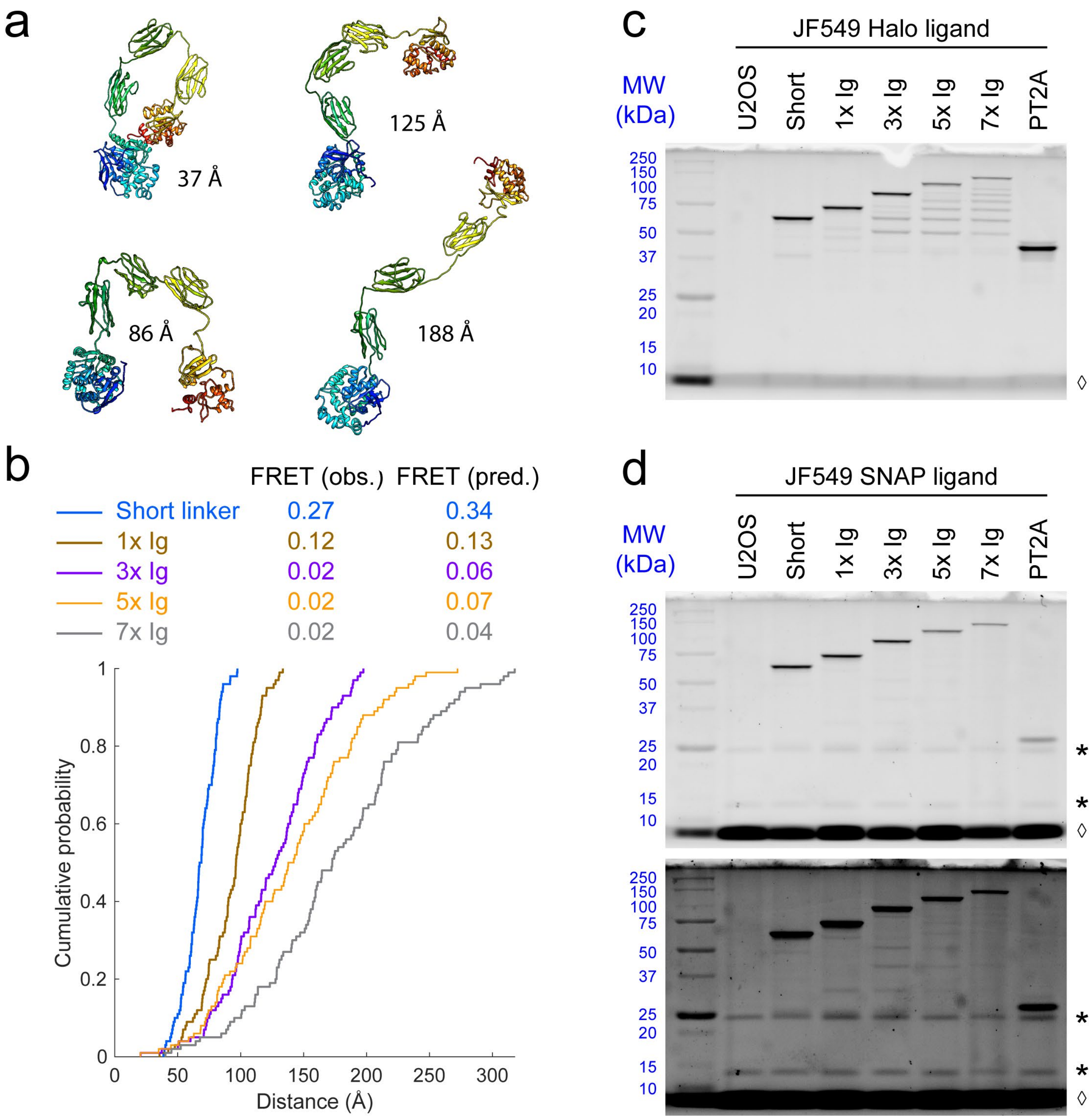

### Supplementary Fig. 4

**SDS-PAGE and PAPA traces of FRB-FKBP. (Related to Fig. 3)**

a) Fluorescence image of an SDS-PAGE gel of lysates from a clonal stable U2OS cell line expressing FKBP-SNAPf and FRB-Halo. Amount of lysate loaded in each lane corresponds to approximately 60,000 cells labeled with either JFX650-STL or JFX650-HTL. ◊, unbound dye.

b) Total fluorescence of the JFX650 receiver fluorophore as a function of frame number when illuminated with red light alternating with pulses of violet light to induce DR and green light to induce PAPA (vertical lines with lightning bolts). The top panel shows the average of cells imaged prior to rapamycin addition (n = 9), and the bottom panel shows the average of cells imaged between 5 and 15 min after rapamycin addition (n = 15). Fluorescence decreased initially due to bleaching and shelving of JFX650 and recovered upon fluorophore reactivation by green and violet light pulses. PAPA (green reactivation) was low prior to rapamycin addition (top panel) and increased following rapamycin addition (bottom panel). DR (violet reactivation) was observed both before and after rapamycin addition. AU, arbitrary units. Intensity traces are displayed without background subtraction.

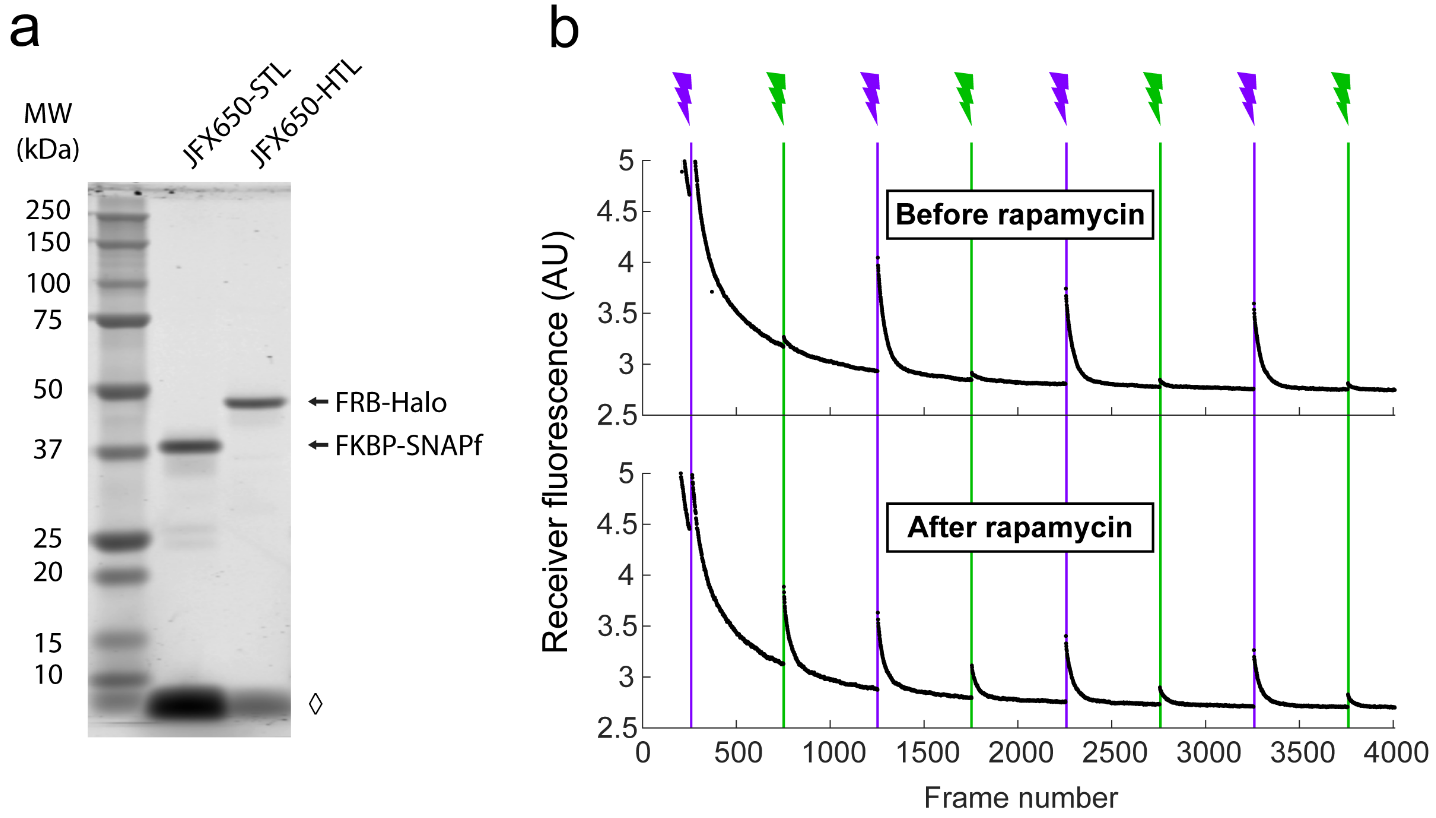

### Supplementary Fig. 5

**SDS-PAGE gels of defined 2-component mixtures and 1-component controls. (Related to Fig. 4 and Supplementary Fig. 6 and 7)**

a-b) Fluorescent SDS-PAGE gels of lysates from stable U2OS cell lines expressing defined 2-component mixtures (1-4) and 1-component controls (5-6). Cells were labeled with either JFX650-STL (S) or JFX650-HTL (H) prior to lysis, and each lane was loaded with a volume of lysate corresponding to approximately 60,000 cells. Associated figure panels are listed for each construct. ◊, unbound dye.

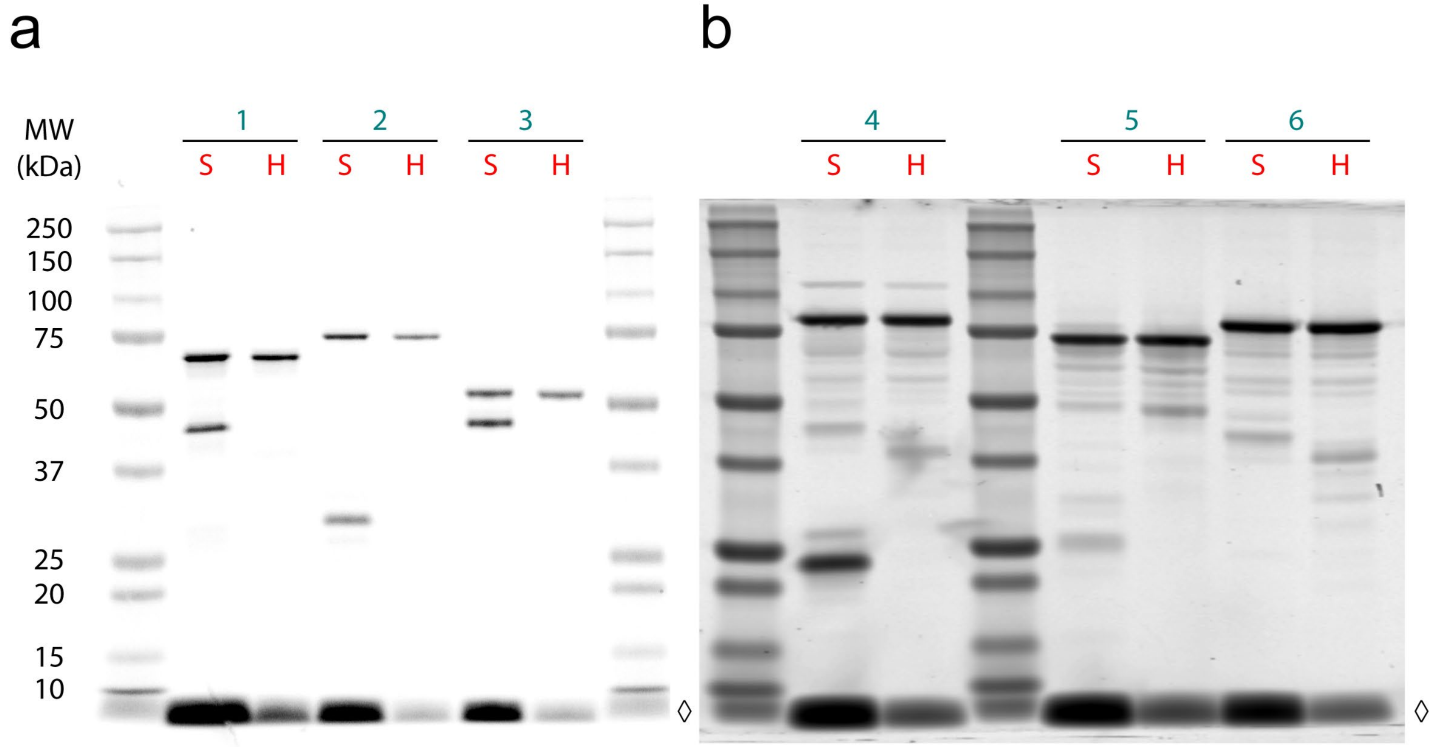


1) Halo-SNAPf-3xNLS-P2A-T2A-H2B-SNAPf-3xNLS (pTG800) Fig. 4a-c, Supplementary Fig. 6a-b
2) SNAPf-3xNLS-P2A-T2A-H2B-Halo-SNAPf (pTG824) Fig. 4d-f, Supplementary Fig. 6c-d
3) SNAPf-60mer-P2A-T2A-Halo-SNAPf (pTG825) Fig. 4g-i, Supplementary Fig. 6e-f
4) Halo-SNAPf-60mer-P2A-T2A-SNAPf (pTG802) Fig. 4j-l, Supplementary Fig. 6g-h
5) H2B-Halo-SNAPf (pTG838) Supplementary Fig. 7d-f
6) Halo-SNAPf-60mer (pTG839) Supplementary Fig. 7g-i
S) JFX650-STL
H) JFX650-HTL

### Supplementary Fig. 6

**Additional analyses of PAPA-SPT experiment with 2-component controls. Related to Fig. 4.**

a,c,e,g) Left panel: Schematic of construct tested. Right panel: A random subset of PAPA (green) and DR (violet) trajectories from an individual cell. Trajectory centroids are aligned to a grid for visualization, and trajectories are displayed in order of increasing average displacement per step. Black scale bar, 1 µm.

b,d,f,h) Histograms of single-frame displacements for single-particle trajectories from all cells. Green, PAPA trajectories. Violet, DR trajectories. The sharp drop in frequency near 1 µm is due to the maximum displacement cutoff used in the particle tracking algorithm.

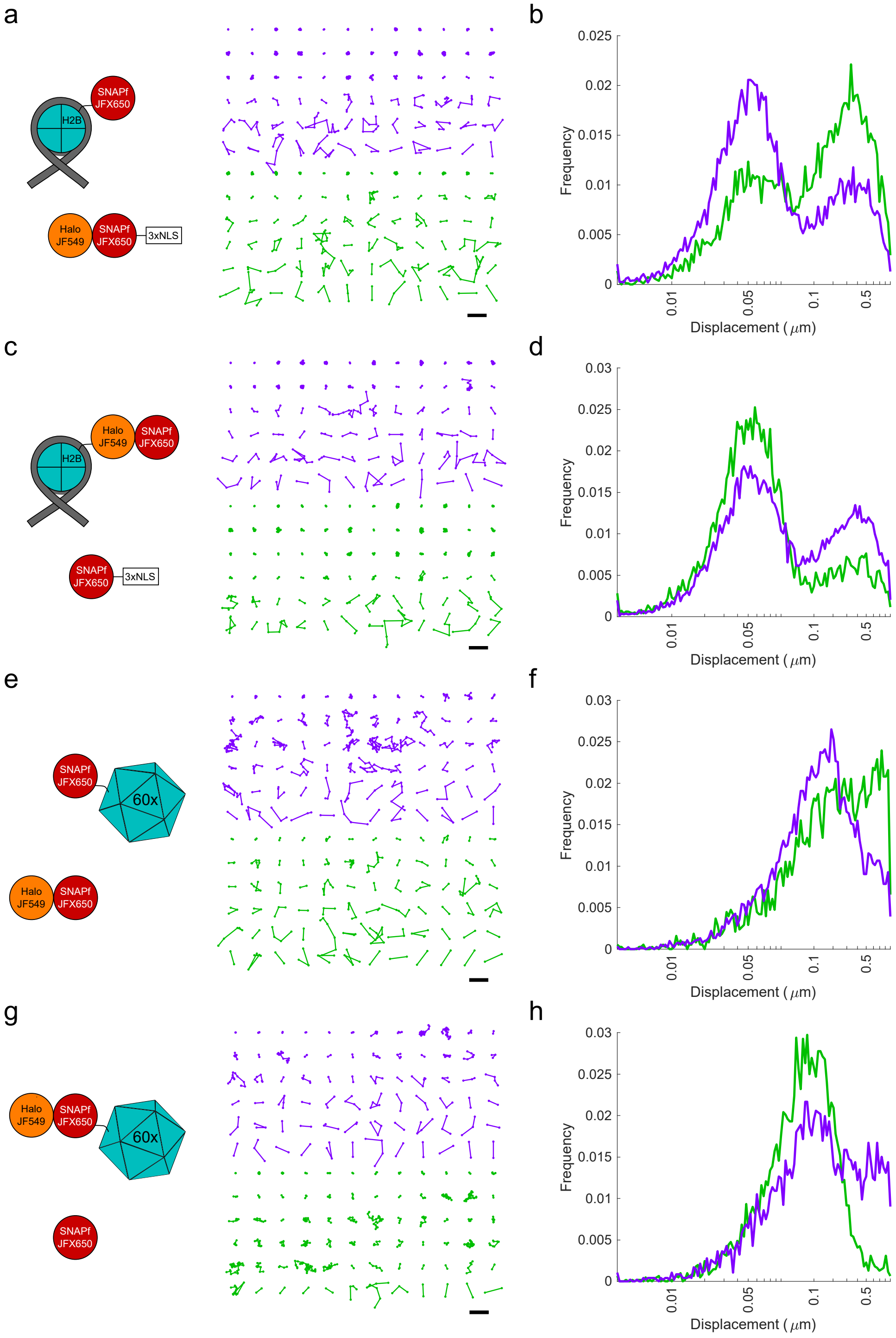

### Supplementary Fig. 7

**PAPA-SPT analysis of single-component controls. Related to Fig. 4.** Individual Halo-SNAPf fusion proteins were labeled with a mixture of JF549-HTL and JFX650-STL, imaged with alternating green and violet photostimulation pulses, and analyzed as in Fig. 4. a-c) Halo-SNAPf-3xNLS. d-f) Halo-SNAPf-H2B. g-i) Halo-SNAPf-60-mer. Note that all 60 subunits are fused to Halo-SNAPf, but only a single label is shown for clarity. b,e,h) Inferred diffusion spectra of PAPA (green-reactivated) and DR (violet-reactivated) trajectories pooled from 10 cells. c,f,i) Fraction bound (c,f) or slow-diffusing (i) among PAPA and DR trajectories from individual cells, obtained from fits to a 2-state (c,f) or 3-state (i) model. P-values calculated using a two-sided paired t-test were 0.58 (c), 0.017 (f), and 0.071 (I).

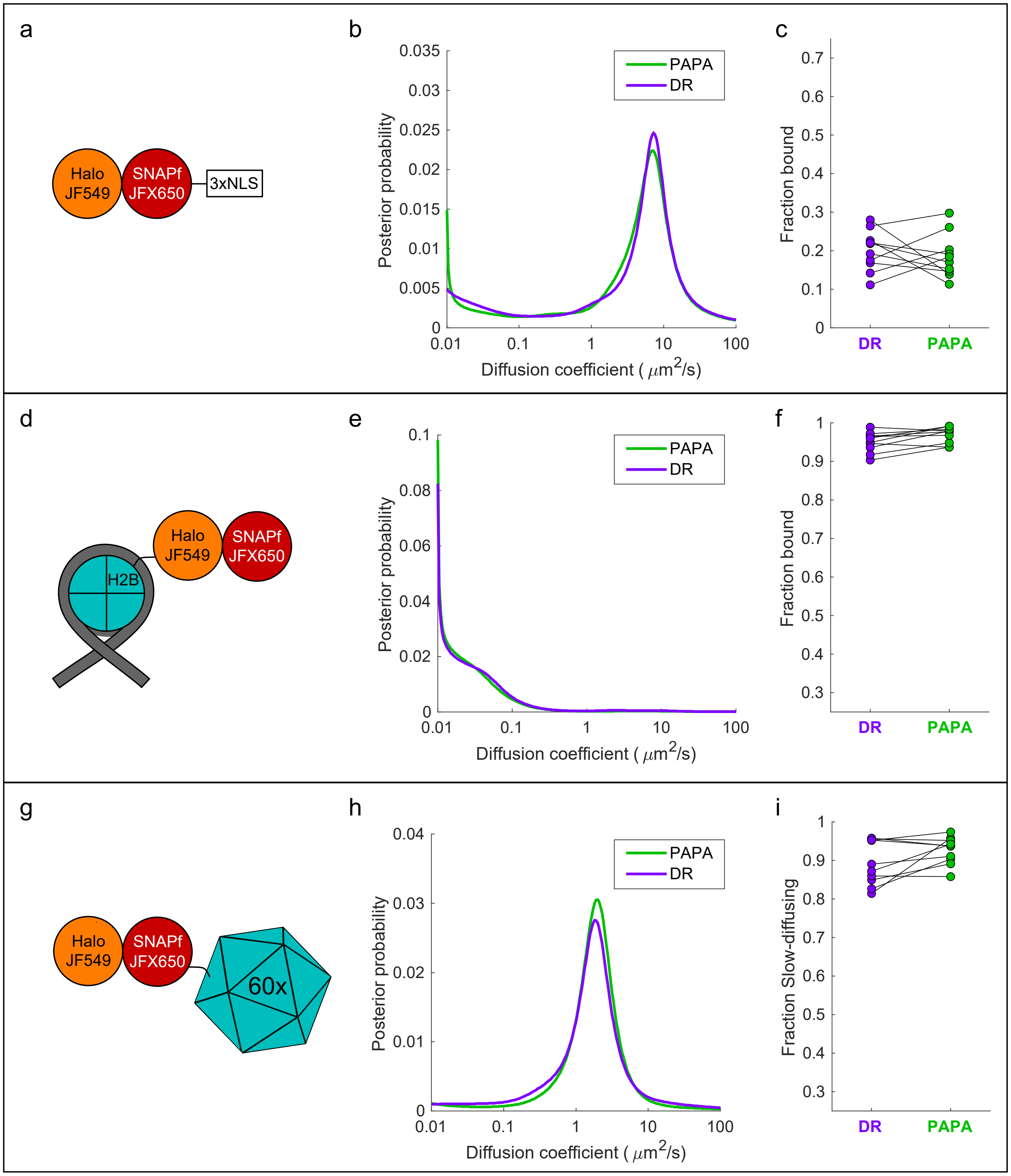

### Supplementary Fig. 8

**PAPA analysis of androgen receptor. Related to Fig. 5.**

a) Fluorescent SDS-PAGE gel of lysates from a clonal stable U2OS cell line expressing SNAPf-mAR and Halo-mAR. Cells were stained with JFX650-STL or JFX650-HTL, and a volume of lysate corresponding to 60,000 cells was loaded per lane. MW, molecular weight in kilodaltons.

b) Ensemble PAPA analysis (see Supplementary Fig. 4b legend) of interaction between SNAPf-mAR and Halo-mAR. PAPA signal (green reactivation) increased after addition of DHT. Intensity traces are displayed without background subtraction.

c) Fraction bound as a function of time relative DHT addition (vertical dashed line) for PAPA trajectories (green) and DR trajectories (violet), based on fits to a 2-state model. Each data point corresponds to PAPA or DR trajectories from a single cell. Solid lines show moving averages over 10-min intervals.

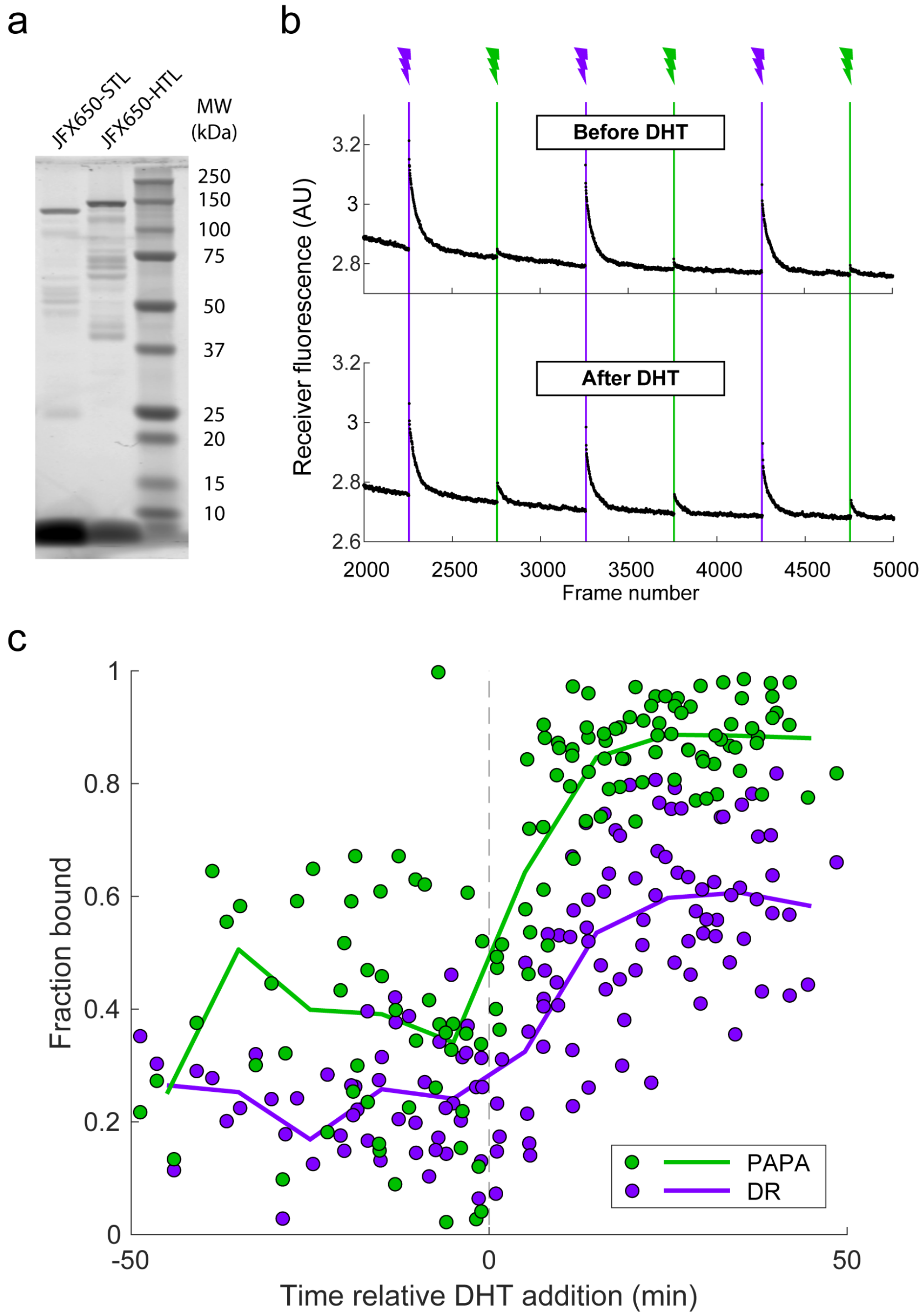

### Supplementary Fig. 9

**Background reactivation by free JF549 dye.** U2OS cells expressing Halo-SNAPf-3xNLS were labeled with JFX650-STL only and incubated with various concentrations of free JF549 dye. PAPA/DR ratio (black points) was measured as in the main text figures by calculating the ratio of reactivation due to 7-ms pulses of 561 nm and 405 nm light. Solid black line shows a linear fit. For comparison, the dashed blue line shows the PAPA/DR ratio measured on the same day for Halo-SNAPf-3xNLS double-labeled with JFX650-STL and JF549-HTL.

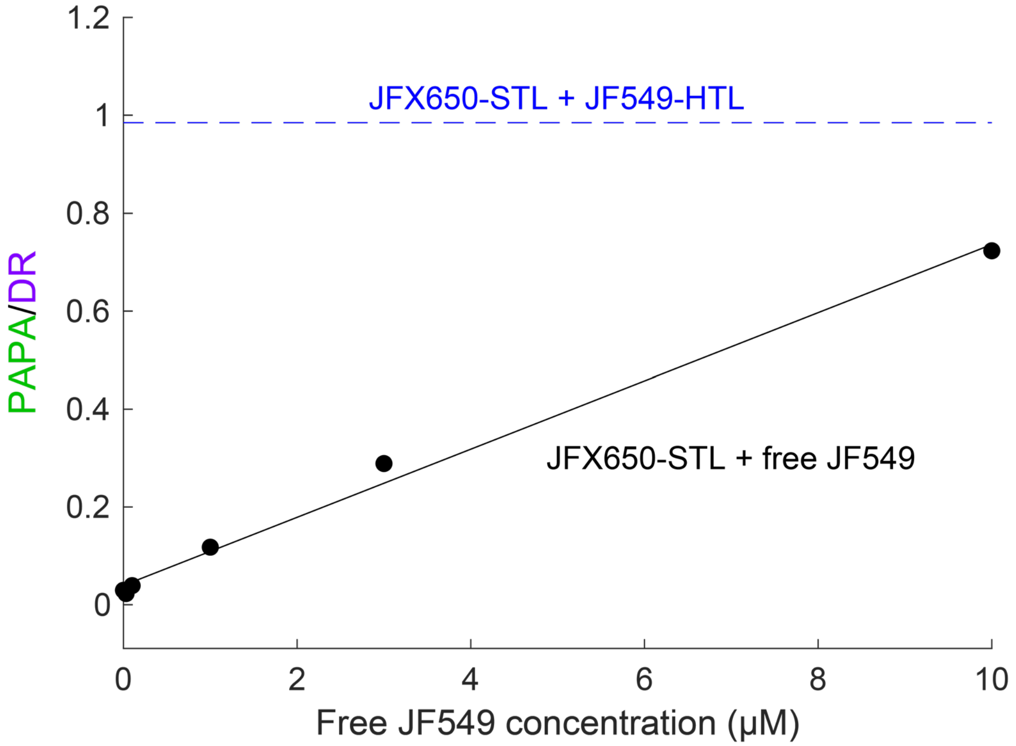

### Supplementary Note 1: SDS-PAGE

Comparison of the SDS-PAGE gels in Supplementary Fig. 3c and 3d shows that the Halo component of the self-cleaving Halo-PT2A-SNAPf fusion is present at a higher concentration than the SNAPf component. This is expected, given that ribosomes often terminate translation at self-cleaving peptide sequences without restarting translation of the next open reading frame (*1*). Although we attempted to flow-sort populations of cells expressing similar levels of Halo, the Halo component of Halo-PT2A-SNAPf is expressed at a somewhat higher level than the other constructs (Supplementary Fig. 3c). Thus, the background PAPA rate observed for Halo-PT2A-SNAPf may overestimate the contribution of nonspecific background to the PAPA rate of the other constructs.

SDS-PAGE analysis of Halo-Ig-SNAPf linker constructs stained with Halo ligand revealed both a full-length protein and a regular ladder of faster-migrating bands, whose molecular weights correspond to Halo fused to different numbers of Ig repeats (Supplementary Fig. 3c). SNAP ligand, in contrast, predominantly labeled the full-length protein (Supplementary Fig. 3d); smaller fragments were only faintly visible with enhanced image contrast (Supplementary Fig. 3d, lower panel). Because the FRET donor (JF549) and PAPA receiver (JFX650) were conjugated to SNAPf, which is almost exclusive to full-length polypeptides, we expect that the presence of smaller Halo-tagged fragments will not impact our measurements of FRET and PAPA efficiency, as these Halo-only fragments will be “invisible” in measurements of JF549-STL fluorescence lifetime and JFX650-STL reactivation.

### Supplementary Note 2: Protein Labeling

In principle, accurate quantification of FRET or PAPA does not require that the FRET donor or PAPA receiver be completely labeled. However, it is critical to thoroughly label the FRET acceptor or PAPA sender, as under-labeling would produce molecules labeled with donor or receiver only (i.e., with no FRET or PAPA), leading to an underestimate of the FRET or PAPA efficiency. Because Halo labeling is much more efficient than SNAPf labeling, we therefore labeled Halo with the acceptor (JFX650-HTL) in FRET experiments and the sender (JF549-HTL) in PAPA experiments, while labeling SNAPf with the opposite fluorophore.

### References

1. Z. Liu, O. Chen, J. B. J. Wall, M. Zheng, Y. Zhou, L. Wang, H. R. Vaseghi, L. Qian, J. Liu, Systematic comparison of 2A peptides for cloning multi-genes in a polycistronic vector. *Sci. Rep.* **7**, 2193 (2017).
